# Object-translocation induces event coding in the hippocampus

**DOI:** 10.1101/2024.06.06.597717

**Authors:** Ya-Li Duan, Shu-Yi Hu, Xiao-Fan Ge, Jia-Li Long, Chu Deng, Yu-Ming Sun, Cheng-Ji Li, Rong Zhang, Xing Cai, Li Lu

## Abstract

Episodic memory refers to the recollection of personally experienced events anchored in spatial contexts. Salient objects within a navigation environment serve as both landmarks and event cues; however, the nuanced integration of these elements during episodic memory formation is not well understood. The mnemonic functions of the hippocampal CA1 region are partially organized along its transverse axis, with proximal and distal subregions preferentially processing spatial and non-spatial information, respectively. Based on comprehensive analysis of neuronal responses along the CA1 proximodistal axis to manipulations of environmental cues and object features, we identified an event-like population coding within CA1 following object displacement. In addition, the neuronal responses at both initial and relocated object locations were shaped by distinct convergent inputs. These findings highlight the specific roles that objects may play in spatial navigation and memory encoding, providing novel insights into the selective processing of object-related information within the entorhinal-hippocampal network.

**Significance Statement:** Salient objects within a navigation environment serve as both landmarks and event cues critical for episodic memory. However, the enigmatic contribution of objects to the “where” and “what” components of episodic memory remains poorly understood. Here we comprehensively analyzed the neuronal responses in the hippocampal CA1 region along its transverse axis, to manipulations of environmental cues and object features. Our results revealed an event-like population coding within CA1 following object displacement, suggesting a minimal landmark role these objects play during memory formation. This study provides novel insights into the selective and flexible processing of object-related information within the entorhinal-hippocampal network.

## Introduction

The hippocampus is crucial for the formation of episodic memory, representing personally experienced events within specific spatial frameworks [1]. In rodents, the collective localized firing of hippocampal neurons, known as place cells, generates a cognitive map of the spatial environment, providing a structural basis for linking episodic memories [2]. Place cells are capable of reorganizing their firing patterns, a process termed remapping, to accommodate environmental changes or novel experiences. This dynamic reorganization facilitates the decorrelation of overlapping neural signals, thereby preventing interference between distinct memories [3, 4]. When objects are introduced into the navigation space, hippocampal place cells undergo substantial changes in activity, including modifications in the size, number, and spatial positioning of their receptive fields [5–10]. Among these diverse responses, some hippocampal cells fire specifically at the location of an object [5], while others fire at a fixed distance and direction from the object, independent of its identity [8]. These unique behaviors, observed in “landmark-vector cells” and “object cells”, likely reflect adaptations to landmark shifts critical for spatial representation and to local object-related events encountered during navigation. This diversity suggests a multifaceted role of salient objects in spatial mapping and memory encoding. However, the complex interplay between landmark and event-related information in the hippocampus complicates the dissection of their individual contributions to the “where” and “what” aspects of episodic memory, with the precise role of objects in these processes still unclear.

The anatomical and functional segregation within the entorhinal-hippocampal network offers critical insights into the complexity of episodic memory formation. The entorhinal cortex (EC) integrates multimodal sensory information and conveys highly processed signals to the hippocampus [11]. Within the EC, the medial region (MEC) contains several specialized cell types, including grid cells [12, 13], aperiodic spatial cells [14], head direction cells [15], border cells [16], object-vector cells [17], and speed cells [18], which collaboratively encode landmark-based distance and directional information, dynamically mapping the spatial location of the animal within the environment [19, 20]. In contrast, the lateral EC (LEC) exhibits minimal spatial selectivity in open fields [21, 22], with neurons predominantly representing non-spatial aspects of the animal’s experience, such as objects [23, 24], odors [25, 26], time [27], memory traces [24], and egocentric positioning relative to landmarks [28], providing the hippocampus with a temporal progression of events [2]. Inputs from the EC are partially segregated along the CA1 transverse axis, where MEC neurons primarily project to the proximal CA1 (pCA1), and LEC neurons preferentially target the distal CA1 (dCA1) [29, 30]. Immediate-early gene expression analyses have indicated distinct mnemonic functions along the CA1 proximodistal axis, with dCA1 preferentially processing non-spatial information, such as time, odor, and objects, across various behavioral tasks, while pCA1 is more engaged in tasks requiring spatial memory [31–36]. Electrophysiological studies have further demonstrated a stronger spatial modulation in pCA1 [37, 38] and a more coherent response to conflicting landmarks in dCA1 [39], as well as coordinated activity between the LEC and dCA1 in odor-associated memory encoding [25, 40]. These findings suggest that the entorhinal-hippocampal network processes landmark and event information with distinct population coding in pCA1 and dCA1, offering a potential resolution to the enigmatic role of objects in self-localization and event integration within episodic memory.

In this study, we aimed to distinguish the “where” and “what” components of local object memory by analyzing place cell activity across the proximodistal CA1 axis as rats navigated an open field with or without objects. Our findings revealed a complex population response to both the objects themselves and the environmental changes resulting from their placement. Manipulating the identity and location of one object elicited stronger population responses in dCA1 than in pCA1, suggesting that the newly introduced objects were primarily processed as events rather than landmarks within the entorhinal-hippocampal network. These results highlight the complexity of information integration that occurs during spatial navigation and memory formation.

## Results

### Object-induced local and remote population responses in CA1

To investigate whether CA1 responses to discrete objects varied along the proximodistal axis, we recorded place cell activity across CA1 subregions (Figure S1) when two identical familiar objects were introduced to a recording box during trials 2 and 3 of a four-trial experimental paradigm (Figure S2A–B). To enable cross-animal comparisons, recording positions were normalized for each tetrode, dividing CA1 into three equal-sized bands: pCA1, middle CA1 (mCA1), and dCA1. While the introduction of objects did not significantly alter overall rat behavior (Figure S2C), CA1 pyramidal cells showed substantial changes in firing patterns, primarily through altered firing rates around the objects (Figure 1A). Notably, dCA1 exhibited the lowest proportion of place cells (Figures S2D & S3) but the highest proportion of rate-modified neurons, while mCA1 contained the highest proportion of stable cells (Figure 1B).

**Figure 1.**
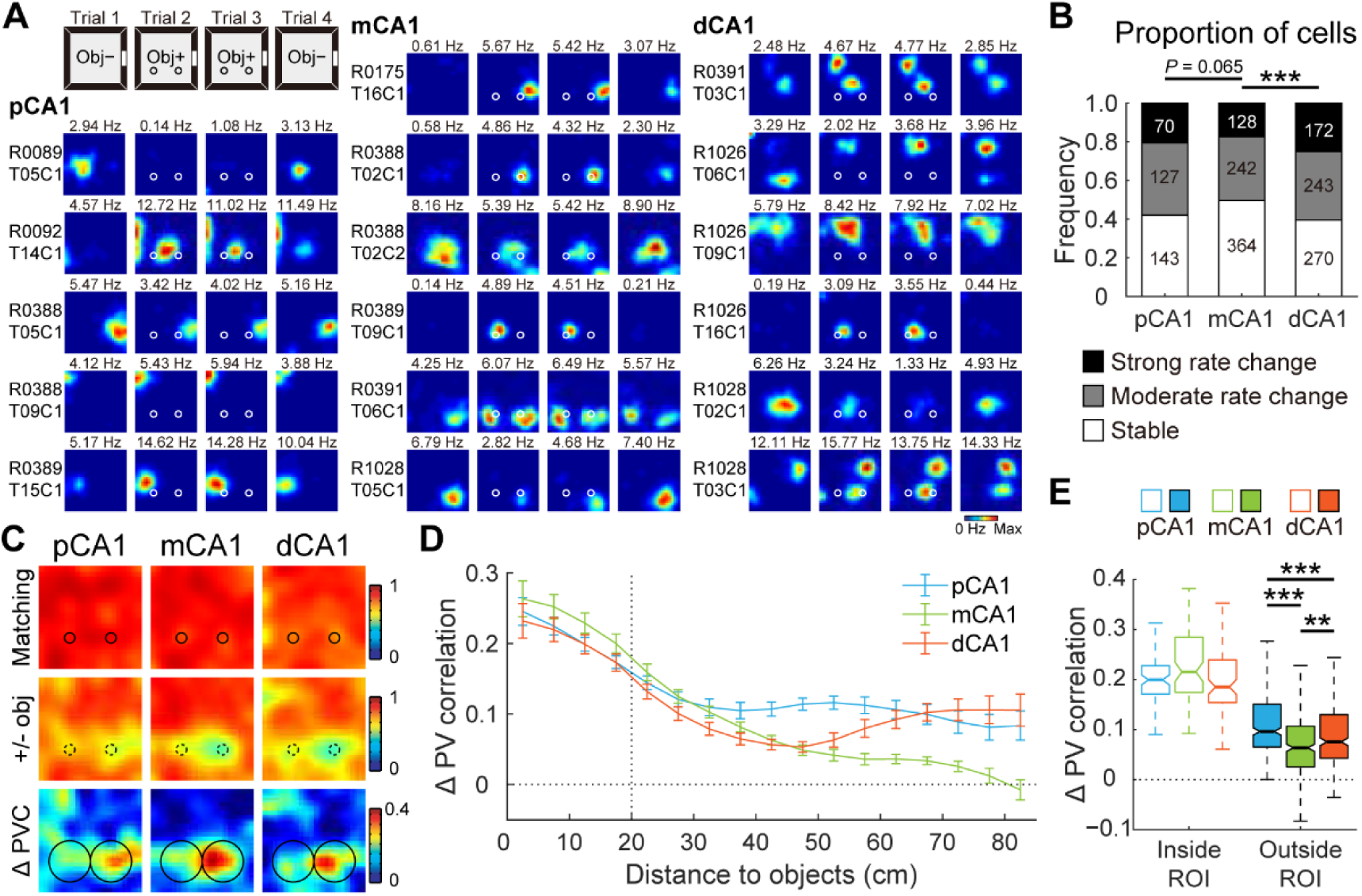
Object-placement induced complex population response in CA1. **A.** Representative firing rate distribution maps (rate maps) of neurons recorded from proximal, middle, and distal bands of CA1, showing responses to familiar objects placed in the recording box. Maps for the same cell across four trials are arranged in rows, with animal and cell IDs indicated on the left. Color is scaled to maximum peak rate of four trials. Text above each map indicates peak rate of that trial. Circles represent objects. Obj−, trial without objects; Obj+, trial with two familiar objects. **B.** Distribution of cells in response to object placement across three CA1 bands. Stable cells had rate changes less than 50%. Moderately remapped cells had 0.5–3-fold increase in rate. Strongly remapped cells had rate changes higher than 3 folds. Numbers indicate active place cells in each category. Proportion of cells with firing rate changes: pCA1, 57.94%, mCA1, 50.41%, and dCA1, 60.58%; chi-square test, χ^2^ = 15.640, *P* < 0.001; Holm-Bonferroni *post hoc* tests, ***: *P* < 0.001. **C.** Heatmaps showing PVC of all putative pyramidal neurons recorded from each CA1 subregion between trials with or without objects (top two rows), as well as changes in PVC between condition pairs (bottom row). Heatmap bins correspond to spatial bins in rate maps. Color scales are indicated on the right of each row. Small circles represent objects, large circles indicate object region. **D.** Degree of changes in PVC as a function of distance to the objects (mean ± SEM) in each CA1 subregion. Two-way ANOVA, distance: F (18, 1 143) = 58.526, *P* < 0.001; band: F (2, 1 143) = 37.619, *P* < 0.001; distance × band: F (36, 1 143) = 5.278, *P* < 0.001. **E.** Boxplots showing degree of changes in PVC between trials pairs, for spatial bins inside and outside the object region, with outliers omitted. Bins inside object region: median (25%–75% percentile), pCA1, 0.199 (0.171–0.228), mCA1, 0.215 (0.174–0.285), and dCA1, 0.185 (0.154–0.239). Bins outside box region: pCA1, 0.096 (0.065–0.150), mCA1, 0.063 (0.026–0.107), dCA1, 0.075 (0.042–0.130); Kruskal-Wallis test, H = 66.740, *P* < 0.001, 960 bins; Holm-Bonferroni *post hoc* tests, **: *P* < 0.01; ***: *P* < 0.001.

Given that hippocampal place cells collectively signal the presence of objects at a population level [10], we analyzed object-related population coding across CA1 subregions using population vector correlation (PVC) to capture composite distribution changes in firing location and rate [41]. Consistent with prior studies [10], trial 4 exhibited a pronounced hysteresis effect (Figure S2E), prompting exclusion of these data from analysis to avoid confounding carryover effects. PVC analysis revealed high correlation between trials 2 and 3 (matching conditions) in all CA1 subregions. As expected, correlations decreased between trials 1 and 2 (before and after object placement), with the most substantial changes in PVC occurring near the objects (Figure 1C). Additionally, dispersed activity shifts distant from the objects were detected in both pCA1 and dCA1, indicating that these responses may not directly reflect the landmark or event features of the objects, but rather environmental changes triggered by object placement. The reduction in PVC gradually diminished with object distance from mCA1, consistent with previous findings [10]; however, pCA1 and dCA1 showed either sustained or increased PVC for regions distant from the objects (Figure 1D). Notably, a remote population response to objects was not observed in the CA3 region, which projects heavily to CA1 (Figure S2F–H), suggesting that direct inputs from the entorhinal cortex may shape this remote response to objects in CA1.

Overall population responses to object introduction differed significantly across CA1 subregions, with the most pronounced PVC reduction observed in pCA1 (Figure S2E&I). To clarify object-induced activity changes, we defined an “object region” as the area within a 20 cm radius around the center of each object, following the distance used in a prior study [10]. While parallel changes in PVC within the object region were observed across CA1 subregions, mCA1 cells displayed minimal activity changes outside the region of interest (ROI) following object placement (Figures 1E and S2J). Thus, the introduction of familiar objects elicited a profound population response in the CA1 region, integrating multiple object-related information components. Neurons in mCA1 demonstrated a comparable local response to the objects but exhibited the weakest remote population reorganization compared to neurons in both pCA1 and dCA1.

### V-shaped population coding of non-spatial information along CA1 transverse axis

To assess whether the remote responses to object placement in pCA1 and dCA1 were indeed driven by changes in non-spatial cues within the arena, we recorded CA1 neuronal activity during a color-reversal task, a conventional non-spatial manipulation paradigm involving the navigation of a square box with alternating black or white walls (Figure S4A–B). As anticipated from prior work [41–43], rate remapping, characterized by alterations in firing rates within place fields while maintaining consistent firing locations [4], was induced in CA1 (Figure 2A). A linear decrease in spatial modulation from pCA1 to dCA1 in the open field was observed (Figures S4–S5), consistent with earlier findings [37, 38]. We first examined population coding of color across CA1 subregions by comparing PVCs between conditions. While population vectors (PVs) correlated strongly between matching conditions in all CA1 subregions, correlations were weaker between black and white wall configurations. Differences in PVC between conditions with reversed wall colors and matching wall colors were significant across subregions (Figures 2B & S6A). Similar to the remote population response to objects, mCA1 exhibited the smallest change in PVC between condition pairs (Figure 2B–C).

**Figure 2.**
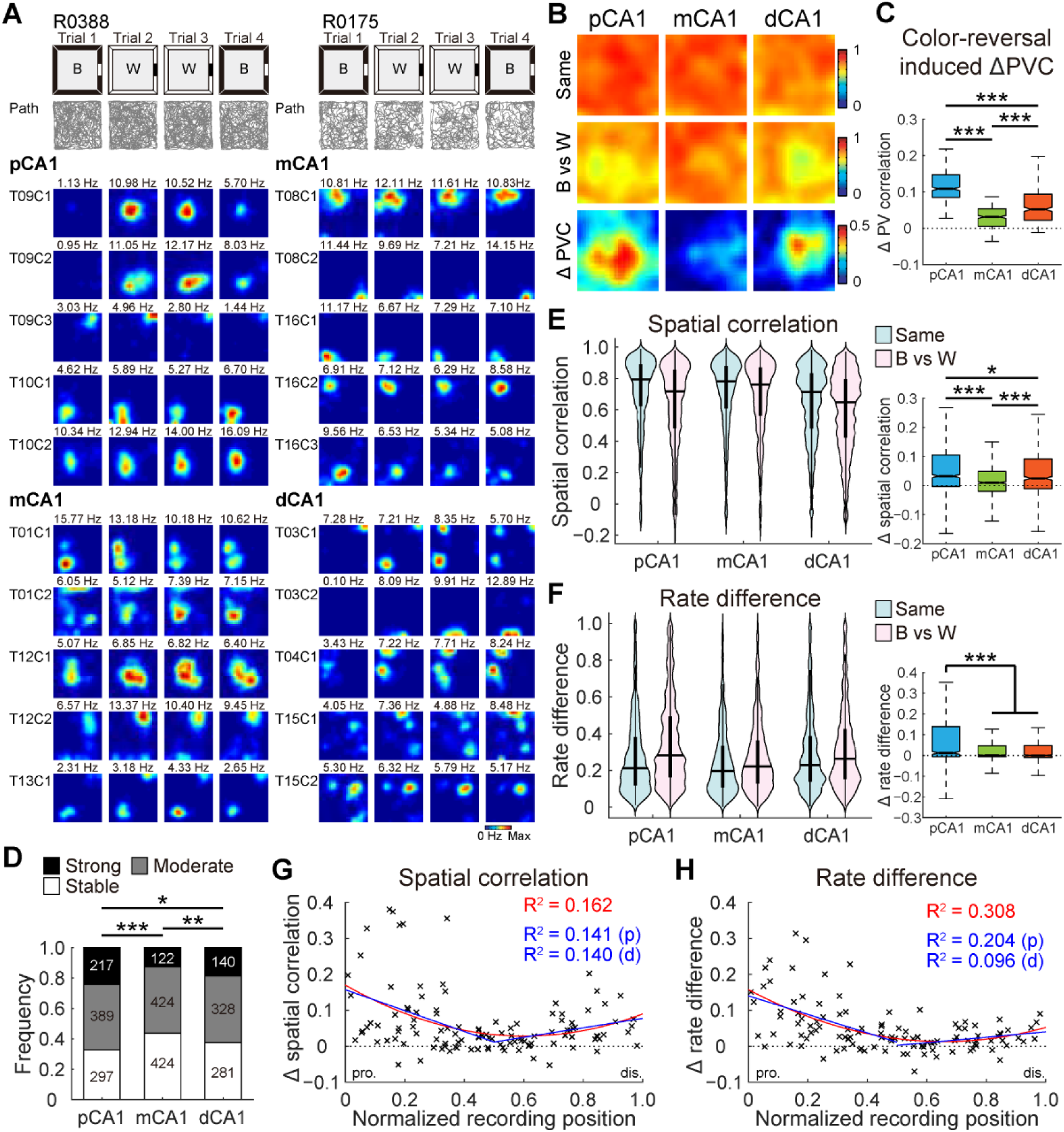
V-shaped population response to reversed wall color across CA1 subdivisions. **A.** Representative rate maps from simultaneously recorded CA1 neurons in two separate color-reversal experiments. Left: pCA1 and mCA1 place cells recorded from rat R0388. Right: mCA1 and dCA1 place cells recorded from rat R0175. Gray lines indicate trajectories of animal in each trial. B, black; W, white version of the same box. Note, mCA1 showed minimum response to the color change. **B.** Heatmaps showing PVC of all putative pyramidal neurons recorded from each CA1 subregion between color conditions (top two rows), and changes in PVC between condition pairs (bottom row) in the color-reversal task. Note, mCA1 exhibited the lowest activity changes. Reduction in PVC is stronger for spatial bins away from arena walls, where the color was manipulated. **C.** Degree of changes in PVC between condition pairs, with outliers omitted. Place cells in mCA1 exhibited least activity change. pCA1: 0.108 (0.085–0.147), mCA1: 0.031 (0.005–0.054), and dCA1: 0.052 (0.022–0.094); Kruskal-Wallis test, H = 541.211, *P* < 0.001, 1 200 bins, Holm-Bonferroni *post hoc* tests, ***: *P* < 0.001. **D.** Distribution of rate remapped neurons between black and white color conditions across three CA1 bands. Stable cells had rate changes less than 50%. Moderately remapped cells had 0.5– 3-fold increase in rate. Strongly remapped cells had rate changes higher than 3 folds. Numbers indicate active place cells in each category. Proportion of strongly remapped cells: pCA1, 24.03%, mCA1, 12.58%, and dCA1, 18.69%, chi-square test, χ^2^ = 41.214, *P* < 0.001; Holm-Bonferroni *post hoc* tests, *: *P* < 0.05, **: *P* < 0.01, ***: *P* < 0.001. **E.** Left: distributions of SC in CA1 subregions between matching (Same) and color reversed (B vs W) conditions. Horizontal line indicates median, and thick vertical bar indicates interquartile range (IQR, 25%–75% percentile) of each subregion. Two-way repeated-measures ANOVA, color: F (1, 2 619) = 262.113, *P* < 0.001; band: F (2, 2 619) = 26.679, *P* < 0.001; color × band: F (2, 2 619) = 28.724, *P* < 0.001. Right: changes in SC between condition pairs, with outliers omitted. pCA1: 0.032 (−0.003–0.106), mCA1: 0.010 (−0.020–0.049), and dCA1: 0.025 (−0.011–0.091); Kruskal-Wallis test, 2 622 cells, H = 68.492, *P* < 0.001; Holm-Bonferroni *post hoc* tests, *: *P* < 0.05; ***: *P* < 0.001. **F.** Similar to **E** but showing changes in RD across CA1 subdivisions. Left: color: F (1, 2 619) = 288.514, *P* < 0.001; band: F (2, 2 619) = 20.373, *P* < 0.001; color × band: F (2, 2 619) = 41.298, *P* < 0.001. Right: pCA1: 0.012 (−0.005–0.140), mCA1: 0.001 (−0.006–0.047), and dCA1: 0.001 (−0.010–0.048); H = 38.540, *P* < 0.001; *post hoc* tests, ***: *P* < 0.001. **G.** Changes in SC between condition pairs as a function of tetrode position. Each cross corresponds to average data of a single tetrode. Blue curve, linear regression; red curve, quadratic polynomial regression. Explained variances (R^2^) are indicated with the same color code. pro., proximal end; dis., distal end. p, proximal half; d, distal half. Proximal half: r = −0.375, *P* = 0.002, Pearson correlation, 63 tetrodes; distal half: r = 0.374, *P* = 0.007, 50 tetrodes. **H.** Similar to **G** but showing correlations between tetrode position and changes in RD between condition pairs. Proximal half: r = −0.452, *P* < 0.001; distal half: r = 0.314, *P* = 0.029.

We next examined rate coding in individual place cells, finding that the proportions of rate-remapped cells between black and white conditions differed across CA1 bands, with mCA1 containing the lowest percentage of rate-remapped neurons (Figure 2D). Rate remapping was quantified using two metrics: spatial correlation (SC), reflecting the degree of rate redistribution for each cell when firing location remains constant, and rate difference (RD), representing the normalized overall change in firing rate irrespective of the firing location of the cell. SC distributions closely resembled PVC distributions along the CA1 transverse axis, with significant SC reductions due to color change in pCA1 and dCA1 compared to mCA1 (Figure 2E). Consistent with the observed SC pattern, RD values also differed significantly between condition pairs and across CA1 subregions, with pCA1 and dCA1 showing more pronounced rate redistribution than mCA1 upon wall color reversal (Figure 2F, left). Notably, pCA1 displayed a significantly higher RD increase between condition pairs (Figure 2F, right). The minimal rate remapping in mCA1 was not due to individual animal variations, as similar patterns emerged when analyses were restricted to simultaneously recorded pCA1, mCA1, and dCA1 neurons (Figure S6B–C). A similar outcome was obtained when the weighted RD for individual place fields (RD*_f_*) was computed (Figure S6D–F).

To visualize population response patterns across the CA1 transverse axis, mean SC and RD values were plotted by tetrode location, revealing a V-shaped distribution from the proximal to distal end of CA1 for both measures. Analyses of proximal and distal tetrodes independently showed significant correlations within each half for SC and RD (Figure 2G–H), with quadratic polynomial curve fitting providing the best explanation for changes along the entire axis. A similar pattern was observed when RD*_f_* was plotted as a function of tetrode location (Figure S6G). To further verify activity change patterns along the CA1 transverse axis, a naïve Bayes decoder was trained on place cell activity from trials with white walls (trials 2 and 3) to predict rat location in a black-walled environment (trial 1). For experiments with over 30 simultaneously recorded place cells across subregions, decoding accuracy was highest in the mCA1 subregion (Figure S6H), indicating minimal activity change between the black and white configurations.

To summary, hippocampal CA1 activity during the color-reversal task demonstrated a V-shaped population response along the transverse axis under non-spatial cue manipulation. Firing rate redistribution was minimum near the middle of the axis and increased progressively towards both the proximal and distal ends. This pattern of population coding closely resembled the remote response observed in the CA1 region to object placement, suggesting that the latter may also be driven by non-spatial environmental changes.

### Extensive reorganization of pCA1 activity in response to landmark changes

Given that the MEC and LEC selectively convey landmark and event-related information to the proximal and distal regions of CA1, respectively [2], we hypothesized that landmark changes would elicit stronger population responses in pCA1, while event changes would lead to elevated activity alterations in dCA1. Thus, we analyzed population activity along the CA1 transverse axis in two rooms with distinct spatial configurations and distal landmarks (Figure S7A–B). Global remapping, characterized by the engagement of a second population of place cells to represent a distinct spatial map in the hippocampus [4], was evident at the individual cell level based on significant changes in the firing location and rate within the recording box (Figure 3A), in line with previous reports [41–43]. PVCs were high for trials within the same room across all subregions but decreased substantially between trials in different rooms (Figures 3B & S7C). The reduction in PVC was region-dependent, with pCA1 exhibiting the greatest decrease, mCA1 showing a moderate decrease, and dCA1 demonstrating the smallest decrease (Figure 3C), suggesting that pCA1 was more sensitive to changes in salient landmarks compared to the distal regions of CA1.

**Figure 3.**
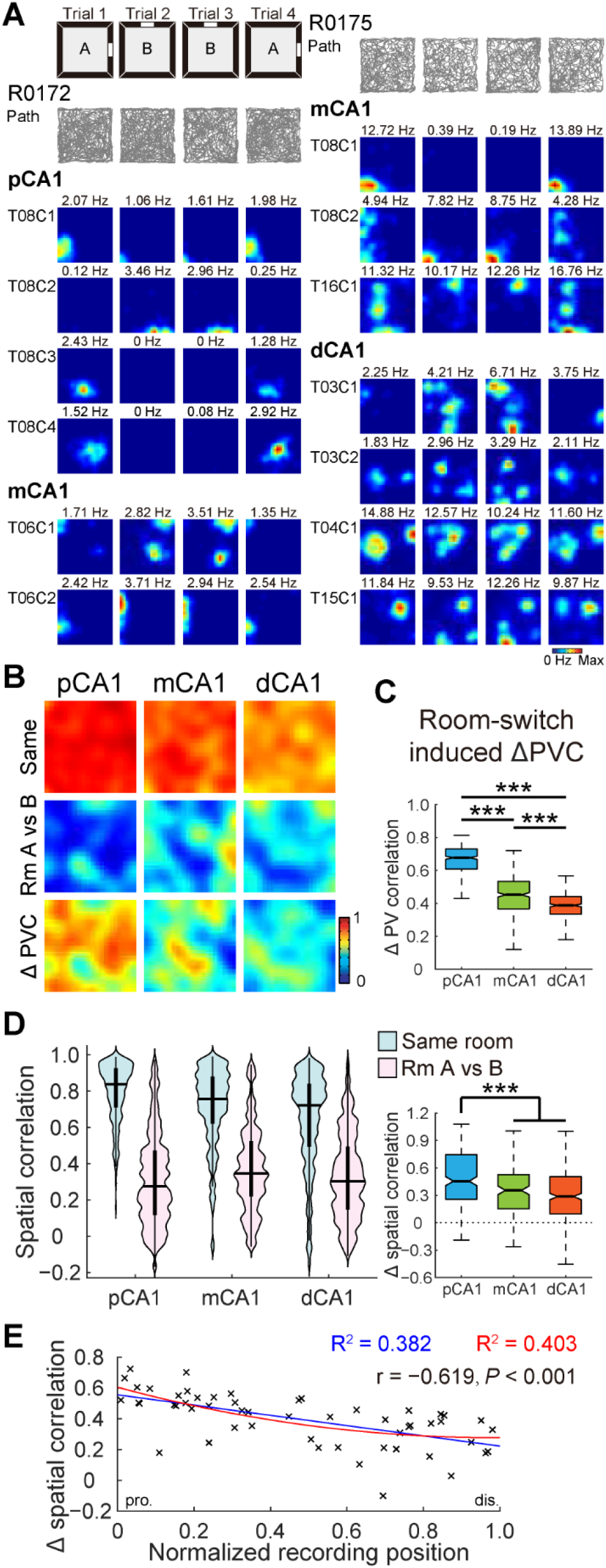
Proximal CA1 was more sensitive to changes in landmarks. **A.** Simultaneously recorded CA1 neurons in two rats exhibiting global remapping in the two-room task. Left: pCA1 and mCA1 place cells recorded from rat R0172. A, Room A; B, Room B. Right: mCA1 and dCA1 place cells recorded from rat R0175. Note, redistribution of firing locations between different rooms. **B.** Heatmaps showing PVCs of all putative pyramidal neurons recorded from each subregion in the same room (top row) and between rooms (middle row), and changes in PVC between room pairs (bottom row). **C.** Degree of changes in PVC between condition pairs, with outliers omitted. pCA1: 0.678 (0.609–0.731), mCA1: 0.452 (0.367–0.535), and dCA1: 0.388 (0.334–0.441); Kruskal-Wallis test, H = 679.871, *P* < 0.001, 1 200 bins, Holm-Bonferroni *post hoc* tests, ***: *P* < 0.001. **D.** Left: distribution of SC in CA1 subdivisions in the same room, and in different rooms (Room A vs B). Two-way repeated-measures ANOVA, room: F (1, 758) = 1 175.415, *P* < 0.001; band: F (2, 758) = 13.538, *P* < 0.001; room × band: F (2, 758) = 27.266, *P* < 0.001. Right: change in SC between condition pairs, with outliers omitted. pCA1, 0.455 (0.256–0.747), mCA1, 0.354 (0.151–0.527), and dCA1, 0.287 (0.095–0.504); Kruskal-Wallis test, 761 cells, H = 41.735, *P* < 0.001; Holm-Bonferroni *post hoc* tests, ***: *P* < 0.001. **E.** Pearson correlation between tetrode position and changes in SC across condition pairs (54 tetrodes). Each cross corresponds to average data of a single tetrode. Blue curve, linear regression; red curve, quadratic polynomial regression.

To further examine spatial sensitivity at the level of individual place cells, we analyzed population overlap across CA1 subregions, which remained consistent (Figure S7D–E). The SC of individual place cells, which is sensitive to rate map reorganization, was high for repeated trials within the same room in all CA1 subregions but decreased markedly between rooms (Figure 3D, left), indicating considerable shifts in firing location across rooms. Between-room SC tended to be higher in mCA1 and dCA1 than in pCA1 (Figure S7F). The decrease in SC with room changes was most pronounced in pCA1 and declined progressively along the proximodistal axis, as reflected by a strong linear correlation between SC decline and tetrode location along CA1 (Figure 3D, right). Quadratic polynomial curve fitting provided minimal additional explanatory power, indicating a predominantly linear gradient from pCA1 to dCA1 (Figure 3E). Similar patterns emerged when using RD to assess neuronal activity levels (Figure S7G–H). Overall, place cell representations were most orthogonalized in pCA1 when salient landmarks were altered, with orthogonality decreasing monotonically along the transverse axis, consistent with previous research [39]. Furthermore, SC variations were significantly correlated with spatial tuning degree (Figure S8), implying a significant contribution of direct MEC inputs to landmark coding within pCA1.

### Enhanced rate coding in dCA1 in response to object replacement

To assess whether the dCA1 region exhibited increased sensitivity to novel events within the environment, we replaced one of the familiar objects with a novel one (Figure S9A), maintaining the landmark properties of the object [17]. Overall, the rats showed consistent behavior before and after object replacement (Figure S9B). A substantial subset of place cells adjusted their firing rates in response to the novel object (Figure 4A), with dCA1 exhibiting the highest proportion of cells responsive to object identity (Figures 4B & S9C). To control for potential hysteresis effects observed in trial 4 (Figure S9D), our analysis focused on firing rate shifts between trials 1 and 2 (object identity switched) and between trials 2 and 3 (object identity unchanged).

**Figure 4.**
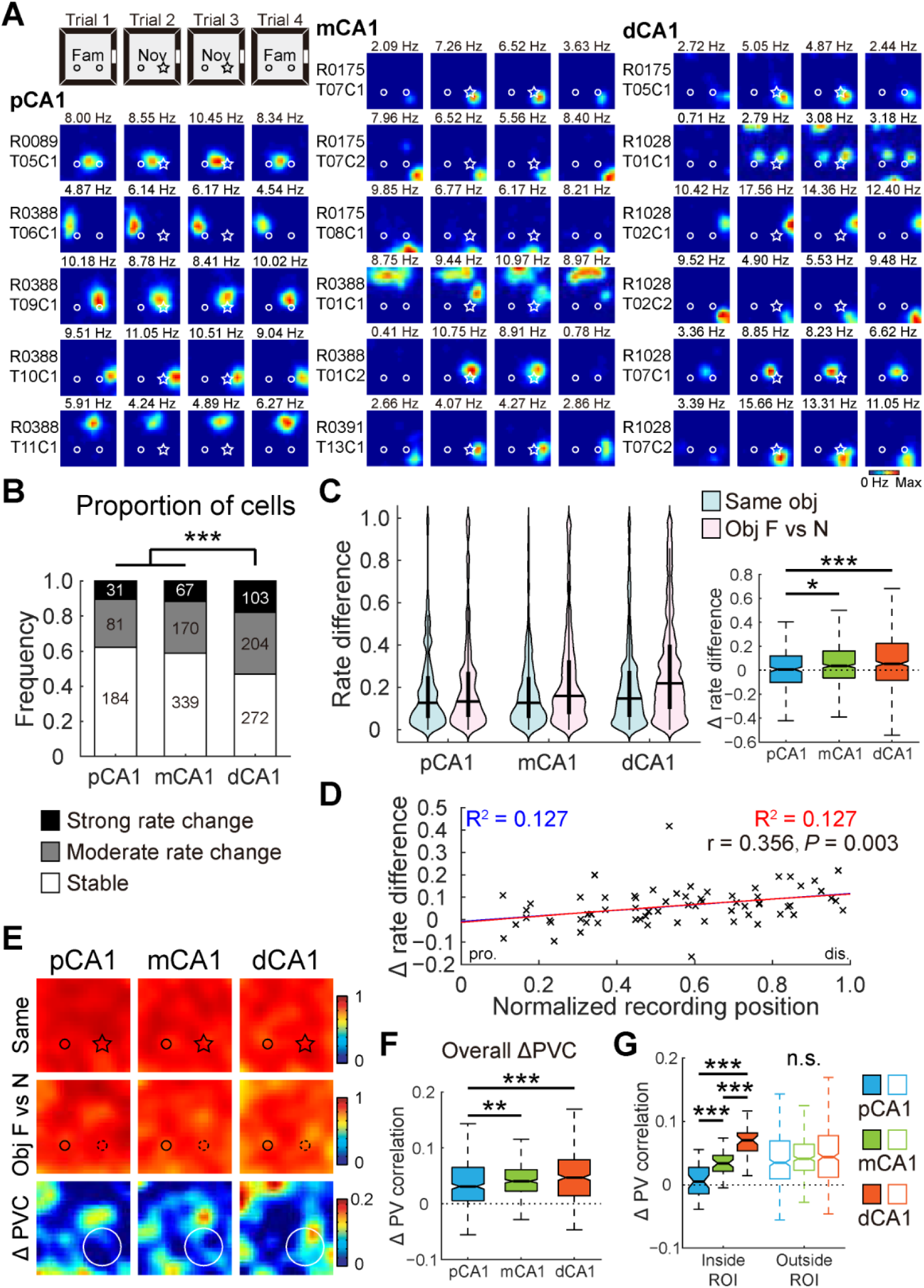
Distal CA1 preferentially represented changes in object-identity. **A.** Representative neurons in proximal, middle, and distal bands of CA1, showing remapping to the novel object. Circle and star represent familiar and novel objects, respectively. Fam, trial with two familiar objects; Nov, trial with right object replaced with a novel object. **B.** Distribution of neurons in response to the novel object across three CA1 bands. Proportion of cells responsive to object-replacement: pCA1, 37.84%, mCA1, 41.15%, and dCA1, 53.02%; chi-square test, χ^2^ = 24.602, *P* < 0.001; Holm-Bonferroni *post hoc* tests, ***: *P* < 0.001. **C.** Left: distributions of RD in CA1 subregions between trials with a novel object (Same) and trials before and after object replacement (Obj F vs N). Two-way repeated-measures ANOVA, novelty F (1, 1 448) = 58.170, *P* < 0.001; band: F (2, 1 448) = 10.527, *P* < 0.001; novelty × band: F (2, 1 448) = 6.169, *P* = 0.002. Right: changes in RD between condition pairs, with outliers omitted. pCA1, 0.008 (−0.101–0.120), mCA1, 0.036 (−0.064–0.162), and dCA1, 0.054 (−0.085–0.225); Kruskal-Wallis test, 1 451 cells, H = 13.604, *P* = 0.001; Holm-Bonferroni *post hoc* tests, *: *P* < 0.05; ***: *P* < 0.001. **D.** Pearson correlation between tetrode position and changes in RD between condition pairs (67 tetrodes). Each cross corresponds to average data of a single tetrode. Blue curve, linear regression; red curve, quadratic polynomial regression. **E.** Heatmaps showing PVCs of all putative pyramidal neurons recorded from each subregion between trials (top two rows), as well as changes in PVC between trial pairs (bottom row) in the object-replacement task. Small circle and star represent familiar and novel objects, respectively; large circle indicates region of the novel object. **F.** Degree of changes in overall PVC between trial pairs across CA1 bands, with outliers omitted. pCA1, 0.031 (0.006–0.065), mCA1, 0.040 (0.023–0.061), and dCA1, 0.047 (0.014–0.079); Kruskal-Wallis test, H = 19.995, *P* < 0.001, 1,200 bins; Holm-Bonferroni *post hoc* tests, **: *P* ≤ 0.01; ***: *P* ≤ 0.001. **G.** Degree of changes in PVC inside and outside ROI between trial pairs across CA1 bands, with outliers omitted. Bins inside ROI: pCA1, 0.005 (−0.014–0.028), mCA1, 0.034 (0.021–0.046), and dCA1, 0.070 (0.053–0.082); Kruskal-Wallis test, H = 61.612, *P* < 0.001, 120 bins. Bins outside ROI: pCA1, 0.035 (0.009–0.069), mCA1, 0.041 (0.023–0.064), and dCA1, 0.044 (0.012–0.078); H = 5.689, *P* = 0.058, 1 080 bins. Holm-Bonferroni *post hoc* tests, n.s. not significant; **: *P* ≤ 0.01; ***: *P* ≤ 0.001.

Due to minimal changes in activity patterns, no significant regional differences in SC changes were detected across trials in the novel-object task (Figure S9E). The overall changes in firing rate for individual place cells were marginally but significantly enhanced in trial pairs before and after object replacement, with the degree of enhancement strongly dependent on the location of place cells. Neurons in pCA1 exhibited the lowest increase in RD, followed by mCA1 and dCA1 (Figure 4C). RD enhancements were significantly correlated with recording positions (Figure 4D). The linear changes in RD were not related to sample size (Figure S9F) or exploratory behavior around objects (Figure S9G). Consistent patterns were obtained for RD*_f_* scores calculated for individual place fields (Figure S9H–I).

The ensemble activity of all pyramidal neurons, as measured by PVC, demonstrated a marginal decrease following object replacement, with a significant interaction between novelty and CA1 subregions (Figures 4E & S9D). Cell assemblies within pCA1 exhibited the smallest response to the novel object (Figure 4F), while Bayesian decoding showed that context prediction accuracy was higher for cells in mCA1 than in dCA1 (Figure S9J). Unlike the object-placement task, population activity reorganization in response to the novel object was less pronounced and less localized across CA1 subregions (Figure 4E). The regional differences in response were primarily attributed to subtle differences in activity changes around the object, with ensemble activity remaining stable outside the object region (Figures 4G & S9K). In addition, CA3 showed minimal response to object replacement (Figure S9L–M). In conclusion, place cells in dCA1 displayed greater firing rate redistribution following object replacement, likely due to preferential input from object-identity-responsive cells in the LEC and potentially from the perirhinal cortex (PER) [22, 23, 44, 45].

### Object-translocation elicited event-like population coding in CA1

Based on the differential encoding of landmarks and events in pCA1 and dCA1 regions, we next examined whether a relocated familiar object within a known environment would contribute to spatial mapping or be encoded as an event. To test this, we translocated a familiar object to a different quadrant of the recording box (Figure S10A). If the object served as a landmark, we anticipated increased activity in pCA1; alternatively, if it functioned as an event, a stronger population response would be expected in dCA1. Despite variations in individual responses to object translocation, the overall discrimination index remained largely unchanged (Figure S10B). A substantial subset of CA1 place cells adjusted their firing patterns, with many neurons displaying new firing fields around the relocated object (current-location coding, CLC), or near its original position (original-location coding, OLC), reminiscent of object-trace cells reported in the LEC [24] (Figure 5A). Notably, although the dCA1 region had the lowest proportion of place cells (Figure S10C), it contained the highest proportion of OLC and CLC neurons, as well as neurons with emergent or disappearing fields outside the region around the relocated object (ROI) (Figure 5B). A minority of cells maintained consistent firing at a fixed distance and direction relative to the relocated object, with or without trace activity remaining in its initial location after relocation, similar to previously reported object-vector cells (OVCs) in the MEC [17]. However, these OVC-like neurons were rare (<2.2% for all subregions) and did not differ across CA1 subregions (Figure 5B).

**Figure 5.**
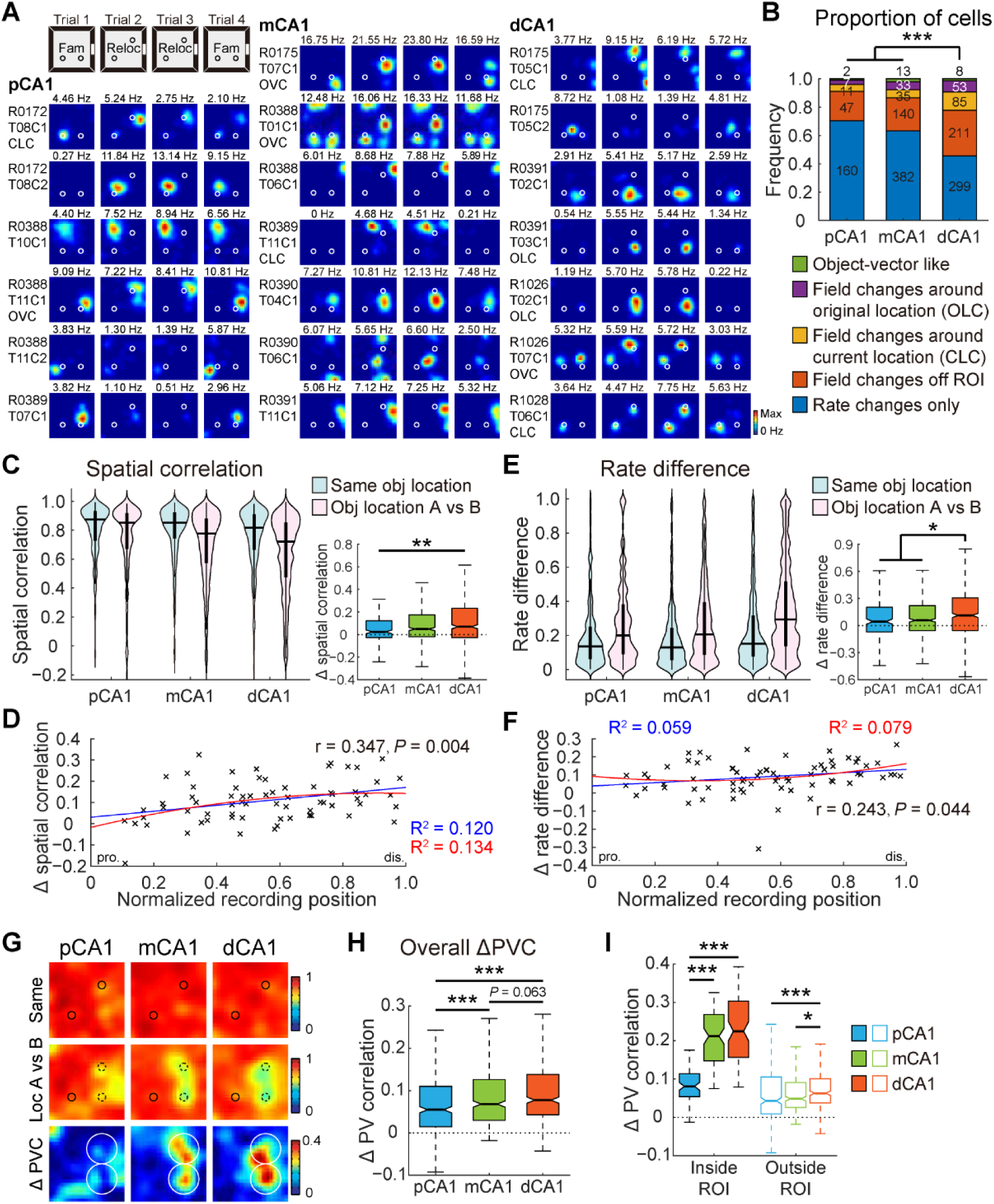
Object-translocation induced stronger population response in dCA1. **A.** Representative neurons in proximal, middle, and distal bands of CA1, showing responses to object displacement. Circles represent objects. Fam, objects in familiar locations; Reloc, trial with right object relocated. **B.** Distribution of neurons in response to object translocation across three CA1 bands. Proportion of cells that showed field changes to the translocation: pCA1, 29.52%, mCA1, 36.65%, and dCA1, 54.42%; chi-square test, χ^2^ = 61.433, *P* < 0.001; Holm-Bonferroni *post hoc* tests, ***: *P* < 0.001. Subgroups: OVC like neurons, Fisher’s exact test, *P* = 0.338; other relocation responsive cells, chi-square test, all *P* < 0.05. **C.** Left: distribution of SC in CA1 subregions between trials with relocated objects (Same obj location) and trials before and after object relocation (Obj location A vs B). Two-way repeated-measures ANOVA, relocation: F (1, 1 483) = 153.986, *P* < 0.001; band: F (2, 1 483) = 21.400, *P* < 0.001; relocation × band: F (2, 1 483) = 4.917, *P* = 0.007. Right: degree of changes in SC between condition pairs, with outliers omitted. pCA1, 0.024 (−0.026–0.121), mCA1, 0.049 (−0.021–0.175), and dCA1, 0.069 (−0.027–0.233); Kruskal-Wallis test, 1 486 cells, H = 10.123, *P* = 0.006; Holm-Bonferroni *post hoc* tests, **: *P* < 0.01. **D.** Pearson correlation between recording position and changes in SC between condition pairs (69 tetrodes). Each cross corresponds to average data of a single tetrode. Blue curve, linear regression; red curve, quadratic polynomial regression. **E.** Similar to **C** but showing changes in RD across CA1 subdivisions. Left: relocation: F (1, 1 483) = 147.858, *P* < 0.001; band: F (2, 1 483) = 20.702, *P* < 0.001; relocation × band: F (2, 1 483) = 2.609, *P* = 0.074. Right: pCA1, 0.044 (−0.073–0.205), mCA1, 0.057 (−0.057–0.221), and dCA1, 0.112 (−0.056–0.308); H = 9.346, *P* = 0.009; *post hoc* tests, *: *P* < 0.05. **F.** Similar to **D** but showing correlations between tetrode position and changes in RD between condition pairs. **G.** Heatmaps showing PVCs of all putative pyramidal neurons recorded from each subregion between trials (top two rows), as well as changes in PVC between trial pairs (bottom row) in object-relocation task. Small circles represent objects, large circles indicate target region. **H.** Degree of changes in overall PVC between trial pairs across CA1 bands, with outliers omitted. pCA1, 0.055 (0.014–0.111), mCA1, 0.068 (0.030–0.127), dCA1, 0.077 (0.043–0.138); Kruskal-Wallis test, H = 44.493, *P* < 0.001, 1 200 bins; Holm-Bonferroni *post hoc* tests, ***: *P* ≤ 0.001. **I.** Degree of changes in PVC inside and outside ROI between trial pairs across CA1 bands, with outliers omitted. Bins inside ROI: pCA1, 0.081 (0.054–0.114), mCA1, 0.212 (0.147–0.269), and dCA1, 0.225 (0.155–0.303); Kruskal-Wallis test, H = 125.736, *P* < 0.001, 240 bins. Bins outside ROI: pCA1, 0.043 (0.009–0.106), mCA1, 0.049 (0.026–0.091), and dCA1, 0.063 (0.038–0.101); H = 17.077, *P* < 0.001, 960 bins. Holm-Bonferroni *post hoc* tests, *: *P* ≤ 0.05; ***: *P* ≤ 0.001.

As anticipated from the distribution of relocation-responsive cell types, the dCA1 region displayed a stronger population response to object relocation compared to its proximal counterpart. The SC of individual place cells was high when objects were in the same location (trials 2 vs 3) but slightly reduced after object relocation (trials 1 vs 2) across all CA1 subregions. The reduction in SC varied significantly by subregion, with the smallest decrease in pCA1 and the largest decrease in dCA1 (Figure 5C). A significant correlation was observed between SC change and tetrode location (Figure 5D). The increase in RD between trial pairs mirrored the SC decrease across CA1 subregions and increased progressively from pCA1 to dCA1 (Figure 5E). A significant correlation was also found between the shift in RD for individual tetrodes and their recording positions (Figure 5F). These spatially graded changes along the CA1 transverse axis were not influenced by object exploration (Figure S10D), sample size (Figure S10E–F), or inter-animal variations (Figure S10G). Consistent findings were observed when RD*_f_* scores were calculated for individual place fields (Figure S10H–I). Correlations between corresponding PVs across spatial bins displayed similar changes to those observed in SC (Figures 5G & S10J). Notably, the decline in PVC between trial pairs progressively increased along the CA1 transverse axis, from pCA1 to dCA1 (Figure 5H). Bayesian decoding also revealed a marked improvement in position predictions within proximally located CA1 regions (Figure S10K). Furthermore, significant reductions in PVC were detected for bins within the ROI in the distal two-thirds of CA1, while pCA1 remained minimally unaffected (Figures 5I & S10L). Collectively, the minimal reorganization in pCA1 alongside the pronounced response in dCA1 suggests that relocated objects were not assimilated into the spatial map as fixed landmarks but were instead encoded as distinct events encountered during navigation.

### Distinct inputs shape responses at original and relocated object locations

Given that object relocation primarily elicited event coding within the CA1 region, we next examined the contributions of specific neuronal subpopulations to this process by systematically removing each subgroup from the overall population analysis. Excluding OVC-like neurons, which were more responsive to landmarks, led to a graded increase in population response to object relocation (Figure 6A). The decline in PVC around the relocated object before and after relocation increased progressively from pCA1 to dCA1 (Figure 6B). To assess differential processing at the original and current object locations, we analyzed population responses at these positions separately and found patterns consistent with those observed within the combined ROI data (Figure 6C).

**Figure 6.**
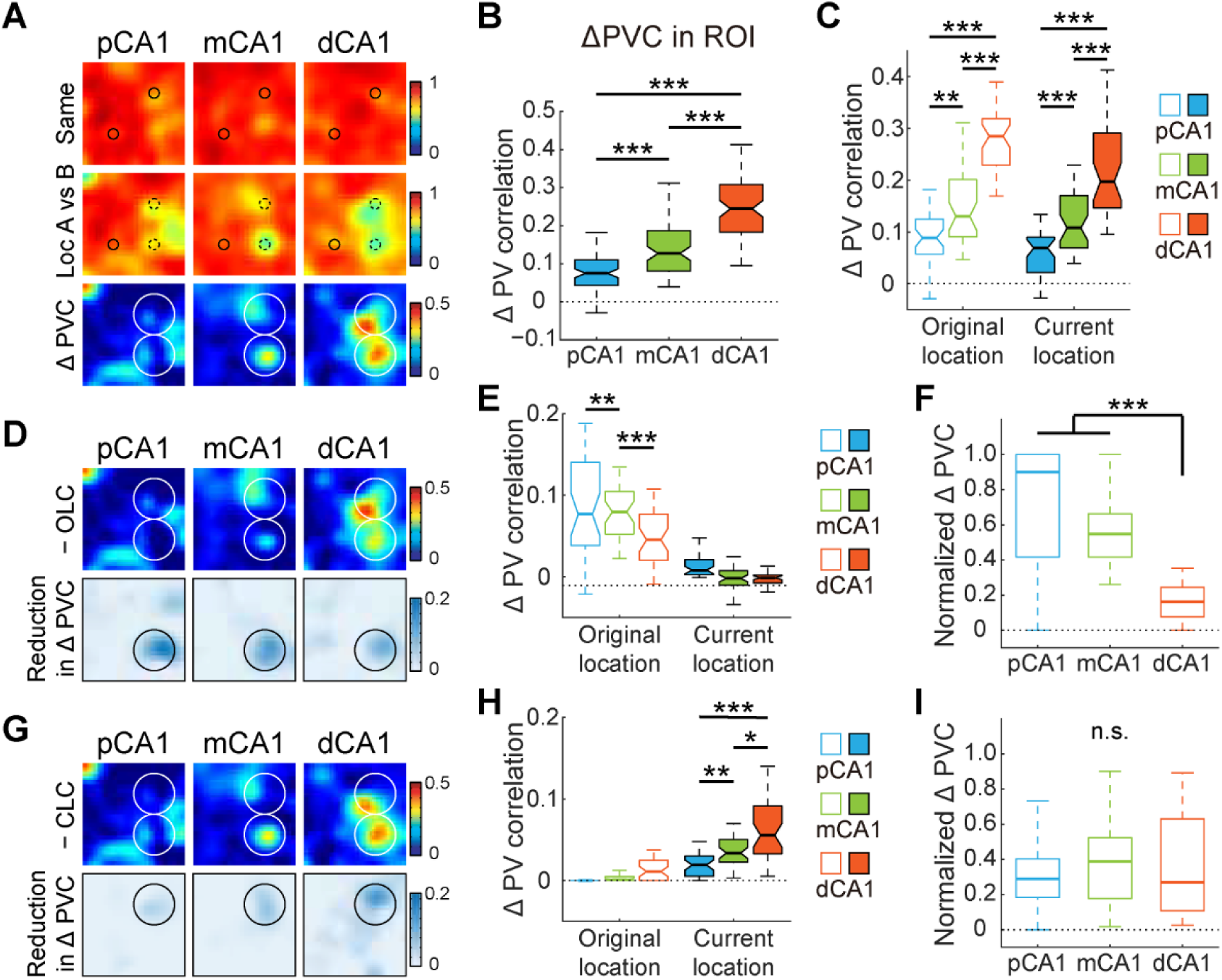
Distinct contributions of neuron subpopulations during event coding. **A.** Heatmaps showing PVCs of place cells excluding OVC-like neurons recorded from each subregion between trials (top two rows), as well as changes in PVC between trial pairs (bottom row) in object-relocation task. Small circles represent objects, large circles indicate target region. **B.** Degree of changes in PVC within ROI between trial pairs across CA1 bands, with outliers omitted. pCA1, 0.075 (0.043–0.110), mCA1, 0.127 (0.081–0.187), dCA1, 0.244 (0.183–0.308); Kruskal-Wallis test, H = 128.394, *P* < 0.001, 240 bins; Holm-Bonferroni *post hoc* tests, ***: *P* < 0.001. **C.** Degree of changes in PVC at original and current locations between trial pairs across CA1 bands, with outliers omitted. Original location: pCA1, 0.089 (0.058–0.125), mCA1, 0.130 (0.091–0.201), and dCA1, 0.285 (0.229–0.319); Kruskal-Wallis test, H = 71.317, *P* < 0.001, 120 bins. Current location: pCA1, 0.069 (0.022–0.090), mCA1, 0.108 (0.069–0.170), and dCA1, 0.198 (0.147–0.291); H = 64.233, *P* < 0.001. Holm-Bonferroni *post hoc* tests, **: *P* < 0.01; ***: *P* ≤ 0.001. **D.** Heatmaps showing changes in PVC for population of place cells without OLC neurons between trial pairs, as well as reductions in PVC change (compared to Fig. 6A bottom panel) after exclusion for CA1 subregions. Circles indicate ROI. **E.** Reductions in PVC at original and current locations after exclusion of OLC neurons across CA1 bands, with outliers omitted. Original location: pCA1, 0.077 (0.038–0.141), mCA1, 0.080 (0.052–0.105), and dCA1, 0.045 (0.020–0.077); Kruskal-Wallis test, H = 15.605, *P* < 0.001, 120 bins. Current location: pCA1, 0.008 (0.003–0.021), mCA1, −0.002 (−0.010–0.008), and dCA1, −0.001 (−0.008–0.002). Holm-Bonferroni *post hoc* tests, **: *P* < 0.01; ***: *P* < 0.001. **F.** Normalized PVC change after OLC neuron exclusion at original location across CA1 bands, with outliers omitted. pCA1, 0.901 (0.417–1.000), mCA1, 0.547 (0.416–0.622), and dCA1, 0.163 (0.077–0.245); Kruskal-Wallis test, H = 71.904, *P* < 0.001, 120 bins. Holm-Bonferroni *post hoc* tests, ***: *P* < 0.001. **G–I.** Similar to **D**–**F**, but show changes in PVC for population of place cells without CLC neurons (**G**), reductions in PVC (**H**. OL: pCA1, −0.0003 (−0.0007–0.0002), mCA1, 0.001 (−0.003–0.005), and dCA1, 0.011 (−0.001–0.025). CL: pCA1, 0.021 (0.010–0.031), mCA1, 0.034 (0.023–0.050), and dCA1, 0.056 (0.033–0.092); Kruskal-Wallis test, H = 34.125, *P* < 0.001, 120 bins), and normalized PVC change at current location (**I**. pCA1, 0.290 (0.183–0.404), mCA1, 0.389 (0.178–0.523), and dCA1, 0.270 (0.107–0.630); H = 0.658, *P* = 0.720). Holm-Bonferroni *post hoc* tests, n.s.: not significant; *: *P* < 0.05; **: *P* < 0.01, ***: *P* < 0.001.

Further removal of OLC neurons led to significant changes at original location but not current location (Figure 6D–E). Notably, the population response to object removal at original object location was predominantly driven by a small fraction of OLC neurons in pCA1 (3.08% of cells contributed to 94.61% of activity changes), while a larger fraction of OLC neurons in dCA1 (8.08%) contributed considerably less to activity change (17.44%) (Figure 6F), suggesting the rapid involvement of other neuronal subgroups in representing past experience from pCA1 to dCA1. Conversely, excluding CLC neurons led to only mild reductions in PVC change at the current location (Figure 6G). Although activity changes showed a graded increase (Figure 6H), the relative reduction in population response due to CLC neuron exclusion remained low (28.56%–35.37%) and was consistent along the transverse axis (Figure 6I), suggesting that population coding for the displaced object was stable across CA1. The profound activity change observed in CA3 at current location, coupled with a weaker response at the original location (Figure S11A–B), suggests that distinct population coding at original and current object locations in CA1 may be partially influenced by CA3 inputs. Excluding neurons with field changes outside the ROI minimally affected responses at either original or current object location (Figure S11C–D). Overall, event coding in CA1 appears to be shaped by signals from multiple sources, with CA3 inputs primarily affecting responses at current location and other inputs influencing responses at original location.

## Discussion

The integration of salient objects during episodic memory formation remains a subject of ongoing debate. Here, we performed a comprehensive analysis of population-level responses across the CA1 proximodistal axis to evaluate the encoding of newly introduced objects in the recording environment, alongside traditional spatial and non-spatial manipulations. Results revealed that object placement triggered localized firing rate changes near the objects and elicited population-level responses similar to those elicited by non-spatial environmental changes (Figures 1–2). Furthermore, by altering spatial configuration and object identity, we validated the segregated encoding of landmarks and events in pCA1 and dCA1 (Figures 3–4). Notably, our final experiment indicated that pCA1 neurons exhibited minimal activity changes in response to object translocation, while more distal neurons along the transverse axis displayed diverse responses to the displaced object, suggesting that the object contributed minimally to self-localization within an established spatial map (Figures 5–6). Collectively, these findings suggest that neutral objects, when introduced into a familiar environment, are primarily processed as transient events rather than integrated landmarks for navigation.

### Experience-dependent integration of landmark and event information

Our results challenge the traditional perspective that object-location memory (OLM) relies predominantly on the MEC over the LEC [46]. While impairments in OLM have been linked to extensive lesions in both the MEC and LEC [47], or complete MEC lesions and inactivation of MEC stellate cells [48–50], LEC-specific lesions appear to have no effect on OLM performance [51, 52]. These discrepancies can be explained by the Hebbian plasticity theory, which posits that the formation of a cognitive map with accurate positional information is highly dependent on repeated experience with landmarks [53]. In standard object-location tasks, brief habituation may not allow for the complete development of a cognitive map, leading to the incorporation of later introduced objects as prominent landmarks. In our study, however, objects were introduced after weeks of intensive training, resulting in event coding tied to a well-established spatial map referencing other established landmarks. The capacity for landmark changes to prompt immediate grid cell reorganization in the MEC, one synapse upstream from the hippocampus, may stem from modified firing patterns in entorhinal object-vector cells that convey allocentric vector information about landmarks [17, 54]. However, these MEC-driven changes showed minimal reflection in the CA1 region during our object-relocation experiments (Figure 5), indicating that selective and flexible processing of MEC inputs in the hippocampus may depend on context-specific mechanisms.

The enhanced population response to object relocation observed in dCA1 likely arises from convergent inputs from multiple brain regions. Within the LEC, subpopulations of neurons are tuned to fire at specific object locations or at certain egocentric distances and directions relative to objects [23, 28]. Object-trace cells within the LEC, which fire exclusively at locations previously occupied by an object, may convey past experiences to dCA1 [24]. These object-location-related and experience-dependent activities are evident in the pronounced activity changes observed at both the original and current object locations in dCA1 (Figure 6). Furthermore, the PER, which projects to the LEC and dCA1, contains object-responsive cells similar to those found in the LEC [29, 44, 45]. Additional object-place signals may also originate from subicular dCA1-projecting neurons, which modulate place cell responses to object displacement [55]. Recently identified subicular corner cells [56] may further contribute to this heterogeneity, although their back-projection to dCA1 remains to be determined.

### Direct entorhinal inputs shape rate and global remapping patterns in CA1

Our study revealed a distinct V-shaped rate remapping pattern within CA1 in both the color-reversal and object-placement tasks (Figures 1–2). The pronounced rate redistribution observed in both pCA1 and dCA1 suggests contributions from both the MEC and LEC to hippocampal rate coding. Certain LEC neurons are known to undergo remapping in response to contextual changes in the environment, and lesions in the LEC have been shown to partially impair hippocampal rate coding [22, 42], supporting the hypothesis that the LEC provides modulated non-spatial inputs critical for hippocampal rate remapping [57]. In contrast, MEC grid cells typically maintain stable firing locations within a given spatial context but exhibit reliable rate redistribution in response to non-spatial manipulations [58–60]. Precise modulation of MEC activity has been linked to hippocampal remapping [61–63], indicating that rate redistribution in MEC cells may directly influence place cell firing. The degree of rate remapping along CA1 may therefore reflect the relative dominance of MEC and LEC inputs across the transverse axis. Neurons in pCA1 or dCA1, predominantly receiving inputs from either the MEC or LEC, exhibit pronounced rate remapping, whereas mCA1 neurons, receiving balanced inputs from both the MEC and LEC, display weaker rate remapping due to dynamic interaction between their competing signals.

Our findings further support the established view that spatial tuning and global remapping in the CA1 region exhibit a gradient decrease from the proximal to distal end (Figures 3 & S4, S5, S8) [37–39]. Global remapping in the rat hippocampus is accompanied by a realignment of activity patterns in grid, head direction, and border cells within the MEC [16, 64], which appears to minimize spatial information overlap and potentially enhance distinct spatial representations in pCA1. Conversely, neurons in the LEC, which encode shared non-spatial features, likely maintain their activity levels, contributing to a more generalized ensemble coding in dCA1 [36, 65]. This gradient in spatial selectivity and global remapping in the CA1 region reflects differential MEC and LEC inputs, with pCA1 predominantly receiving spatial signals from the MEC and dCA1 primarily receiving non-spatial signals from the LEC [66].

### Differential representation of contextual cues along CA1 transverse axis

We also identified three distinct cue representation patterns along the proximodistal axis of CA1, which may contribute to hippocampal mnemonic functions. These patterns comprise a gradient of decreasing spatial information processing, a V-shaped response to non-spatial environmental changes, and an increase in the representation of local events. These variations likely stem from selective inputs from specialized neuronal populations in the entorhinal cortex (Figure S12) [17, 19, 22–24, 28]. Our findings suggest the need for further investigation into the causal dynamics linking selective upstream inputs to specific downstream representations. Future theoretical and experimental endeavors should aim to elucidate the precise mechanisms that govern these interactions, providing crucial insights into the neural basis of spatial navigation, episodic memory, and the broader cognitive functions of the hippocampus.

### Limitations of the study

This study has two primary limitations. First, CA1 neuronal activity was recorded using a hyperdrive with movable tetrodes, constraining our capacity to capture activity from a large population of neurons across CA1 subregions simultaneously in individual rats. This restriction may have introduced inter-animal variability, potentially confounding the interpretation of our results. Second, the complexity in CA1 extends to its radial organization into superficial and deep sublayers, each with distinct characteristics and functions [67]. Our tetrode-based methodology was unable to differentiate between these sublayers. As such, future studies should incorporate two-photon calcium imaging and multi-shank probe recording to monitor neural activity within specific sublayers. These advanced techniques could provide further insights into laminar contributions to functional heterogeneity along the CA1 transverse axis, leading to a more nuanced understanding of landmark and/or event coding within the hippocampus.

## Methods

### Subjects

The experiments were conducted at the Kunming Institute of Zoology, Chinese Academy of Sciences (Kunming, China) in compliance with the guidelines for the care and use of laboratory animals. The study received approval from the Institutional Animal Care and Use Committee of the Kunming Institute of Zoology (KIZ-IACUC-RE-2021-06-019). Data were collected from 22 male Long-Evans rats (3–4 months old, 500–600 g at the time of implantation). Five rats from a previous study [43] contributed proximal-to-middle CA1 data. The remaining 17 rats were implanted with a multi-tetrode “hyperdrive” targeting the right hippocampus. The animals were maintained on a reversed 12-hour light/dark cycle (lights off from 9.00 am to 9.00 pm) under standard laboratory conditions (temperature: 20–23 °C, relative humidity: 40%–60%). Following surgery, the rats were housed individually, and experimental procedures were conducted during the dark phase. During the experimental period, the animals were mildly food restricted to maintain their weight at 85%–90% of free-feeding body weight.

### Electrode preparation

The multi-tetrode ‘‘hyperdrive’’, consisting of 18 independently movable tetrodes made of 17 µm polyimide-coated platinum-iridium (90%–10%) wire (California Fine Wires, USA), was prepared as described previously [68]. The electrode tips were plated to reduce impedance to 150–250 kΩ at 1 kHz (Biomega, Bio-Signal Technologies, China) and sterilized with ethylene oxide (Sanqiang Medical, China) within 48 hours before implantation.

### Surgery

Rats were anesthetized with isoflurane (air flow: 0.8–1.0 L/min, 0.5–3% isoflurane vol/vol mixed with oxygen), with the concentration adjusted based on physiological monitoring (RWD Life Science, China). Upon induction of anesthesia, Metacam (2 mg/mL, 1 mg/kg body weight), Baytril (50 mg/mL, 5 mg/kg), and atropine (0.5 mg/mL, 0.1 mg/kg) were administered subcutaneously. The animal’s head was then fixed in a stereotaxic frame (RWD Life Science, China). Local anesthetic (2% lidocaine, 200 μL) was applied under the skin before making the incision. The skull and dura were removed, followed by immediate hyperdrive implantation. Tetrodes were inserted at the cortical surface above the right hippocampus (CA1: anteroposterior (AP) from 3.2 to 5.6 mm, mediolateral (ML) from 1.0 to 4.5 mm relative to bregma; CA3: AP: 4.2 to 4.9 mm, ML: 2.4 to 4.0 mm). The hyperdrive was secured to the skull with jewelers’ screws and dental cement. Grounding was achieved by connecting two screws above the cerebellum to the hyperdrive ground, while two reference tetrodes were positioned in an inactive area of the cortex for differential recording.

### Electrophysiological recordings

For the first 3–4 weeks post-surgery, the tetrodes were advanced daily until they reached the dorsal CA1 pyramidal cell layer, characterized by large-amplitude theta-modulated complex-spike activity. On the recording day, the electrodes were kept stationary to ensure stable recordings. Broadband neural signals were captured from the hyperdrive using a multichannel data acquisition system (Zeus, Bio-Signal Technologies, China). Unit activity was amplified (× 192) and band-pass filtered (300 to 7 500 Hz). Spike waveforms that exceeded the −50 μV threshold were time-stamped and digitized at 30 kHz for 1 ms. Local field potential (LFP) signals, one per tetrode, were recorded in the 0.3–300 Hz frequency band at a sampling rate of 1 000 Hz. The animal’s position was monitored in real-time by tracking light-emitting diodes (LEDs) on the headstage at 50 Hz using a Cyclops video tracking system (Bio-Signal Technologies, China).

### Behavioral paradigms

Each task consisted of four 10-minute exploration trials within the recording box, interspersed with five 5-minute rest periods in a flowerpot placed on a pedestal outside the box. The exploration trials were designed in an A-B-B-A configuration, where A and B represented distinct test conditions. To encourage exploration, cookie crumbs were randomly scattered throughout the recording box. One week after surgery, the mildly food-restricted rats were trained daily in the color-reversal task in two separate rooms until they could traverse the entire arena within 10 minutes in all trials. Once the tetrodes reached the CA1 cell layer, the animal underwent sequential testing and recording in the color-reversal, two-room, and three object-related tasks.

### Test environments

Recording boxes (100 cm × 100 cm) consisted of a metal frame and four individually exchangeable plastic walls (black on one side, white on the other; 100 cm × 50 cm). Cue cards (white cue card for black walls and vice versa; 50 cm × 30 cm) were placed at the center of one wall. Distal background cues were obscured by curtains encircling the recording box (120 cm × 120 cm) for the color-reversal and object-related tasks but not for the two-room task. The pedestal with the flowerpot was positioned between the test box and experimenter, outside the curtains. The recording rooms for the two-room task were similar in size but featured different spatial arrangements.

### Color-reversal task

As detailed previously, this task was designed to elicit rate remapping in the hippocampus [41–43]. Rats were tested in the recording box at a fixed location over four trials daily, with trials 1 and 4 conducted with black walls and trials 2 and 3 conducted with white walls. The floor of the recording box was cleaned with 75% ethanol after each trial.

### Two-room task

This task was designed to evoke global remapping in the hippocampus [41, 43]. Rats were tested in similar recording boxes with black walls located at fixed locations in two separate recording rooms, with trials 1 and 4 conducted in room A and trials 2 and 3 conducted in room B. The floor of the recording box was cleaned with 75% ethanol after each trial.

### Tasks with objects

#### Object-placement task

Animals were tested in the black recording box at a fixed location over four trials, with two identical familiar objects (glass bottles filled with white stones) placed in quadrants 3 and 4 of the box for trials 2 and 3. <u>Novel-object task:</u> Animals were tested in the black recording box containing two objects over four trials, with the familiar object in quadrant 4 replaced by a novel object in trials 2 and 3. <u>Object-relocation task:</u> Animals were tested in the black recording box containing two familiar objects over four trials, with the object in quadrant 4 relocated to quadrant 1 for trials 2 and 3. For each task, the floor of the recording box and the objects were thoroughly cleaned with 75% ethanol before each trial.

### Occupation time and discrimination index

Occupation time (OT) and discrimination index (DI) were employed to quantify preference for objects in the object-placement task and for relocated/novel objects in the object-relocation and novel-object tasks. OT was defined as the proportion of time spent around the objects relative to the total time spent in the recording box, and DI was calculated as:

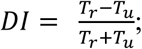

where *T_r_* is the time spent around the relocated/replaced object and *T_u_* is the time spent around the unchanged object. A higher DI value indicates a greater preference for the relocated/novel object.

### Histology and recording sites

Following the completion of electrophysiological recordings, the electrodes were maintained in their positions. The rats were subsequently euthanized and perfused with formalin for brain extraction. Frozen brain sections (40 μm thick) were cut on a cryostat (KEDEE, KD-2950, China), and stained using cresyl-violet (Sigma Aldrich, USA, CAS: 10510-54-0, Cat# C5042-10G). the Nissl-stained brain sections were imaged using a bright-field microscope (2×/0.06, 4 ×/0.13 NA objectives, COIC, UB103i, China), with UOPView software (v2.0). The location of each tetrode tip was then identified and recorded. Hippocampal subregions were delineated based on established criteria [29, 37]. ImageJ (v1.52a) was used to estimate the normalized position along the proximodistal axis of the CA1 region for each recording site. Hand-drawn lines tracing the CA1 pyramidal cell layer were extended from the CA2/CA1 border to the CA1/subiculum border, passing through the recording site on each section. The distance from the CA2/CA1 border to the recording site was measured and normalized by the total length of the CA1 region on the same section. Based on these normalized positions, recording sites were categorized into proximal (pCA1, 0–0.33), middle (mCA1, 0.33–0.67), and distal (dCA1, 0.67– 1) subregions.

### Spike sorting and cell classification

Spike sorting was manually performed using the MClust graphical cluster-cutting tool in Matlab (v4.4, A.D. Redish). The process involved the analysis of two-dimensional projections of the multidimensional parameter space, consisting of waveform amplitudes, waveform energies, and waveform peak-valley differences. Waveform and autocorrelation functions were also employed as tools for separation and classification. Clustering was applied on the entire dataset of each experiment, with the spikes subsequently divided into four exploration trials.

Putative excitatory cells were distinguished from putative interneurons and bypassing axons based on spike width (peak-to-trough waveform durations greater than 250 μs), average rate (less than 5 Hz), and occasional presence of bursts. Only cells classified as putative excitatory neurons were included in subsequent analyses. Cells recorded repeatedly in the same task were excluded from analyses.

### Rate maps and place fields

Position estimates were derived from tracking the LEDs on the headstage connected to the hyperdrive. Data were filtered to include only epochs with instantaneous running speeds between 2.5 and 100 cm/s. The mean firing rate for each cell was calculated by summing the total number of spikes and dividing it by the time spent in that trial. Place field analyses were conducted exclusively for cells with a mean firing rate of 0.10 Hz or higher, defined as active cells. Spatial firing rate distributions (place fields) for each well-isolated neuron were generated by summing the spikes within specific location bins (5 cm × 5 cm), dividing by the duration of time spent by the rat in that location, and applying Gaussian smoothing centered on each bin, as described previously [43]. A place field was defined as a contiguous region of at least 225 cm^2^ (nine or more 5 × 5 cm bins) where the firing rate exceeded 20% of the peak rate and the peak firing rate of the area was 1 Hz or greater. The number of non-overlapping place fields was estimated for each cell.

### Neuron subpopulations in the object-relocation task

Neurons classified as object-vector cell-like were characterized by their exclusive activation at a specific distance and direction relative to the displaced object, both before and after its relocation. Furthermore, the center of the receptive field must be situated within a region 20 cm around the object. Neurons designated as original and current location coding cells were identified by the presence or absence of at least one receptive field at the object’s original or current location (within a 20 cm radius), respectively.

### Theta modulation

The theta modulation index (TMI) was computed to assess the temporal modulation of firing within the theta frequency range (7–10 Hz) of each cell, as described previously [69]. A temporal autocorrelogram was constructed by tallying the occurrences of spikes within each 5-ms bin over a range of 0 to 500 ms from a reference spike at time 0, then dividing by the total trial length to determine the rate of occurrence for each bin. The TMI was calculated as:

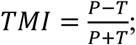

where *P* and *T* represent the theta modulation peak (mean of bins 100–140 ms) and trough (mean of time bins 50–70 ms), respectively.

### Spatial information, coherence, and stability

The spatial information rate in bits per second (*SI_r_*) and spatial information content in bits per spike (*SI_c_*) were calculated as described in previous research [70]:

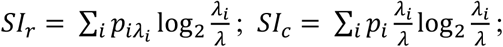

where *λ_i_* is the mean firing rate of a unit in the *i*-th bin, *λ* is the overall mean firing rate, and *p_i_* is the probability of the animal being in the *i*-th bin (occupancy in the *i*-th bin / total recording time). Pyramidal neurons with *SI_c_* scores above the 95^th^ percentile of the shuffled distribution were classified as place cells.

Spatial coherence was estimated as the first-order spatial autocorrelation of the unsmoothed place field map, i.e., mean correlation between the firing rate of each bin and average firing rate in the eight adjacent bins.

Spatial stability was determined by comparing the firing pattern in the first half of each trial to that in the second half. The rate map for each half was smoothed and binned into 5 cm × 5 cm matrices, and the firing rates in the bins of the two half-trial maps were correlated for each trial.

### Spatial correlation (SC)

Firing patterns were compared across trials using a SC procedure. Each map was smoothed and binned into 5 × 5 cm matrices, and the firing rates in the pixels of the two maps were correlated for each cell. SC between two inactive trials (firing rates below 0.1 Hz) was excluded to avoid artifacts. For the two-room task, rate maps were rotated in 90° steps to identify the maximum SC score when compared across rooms.

### Rate difference (RD) and rate difference by field (RD*_f_*)

The change in overall firing rate across trials was calculated as RD:

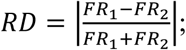

where *FR_1_* is the mean firing rate of a cell in one trial and *FR_2_* is the mean firing rate of the same cell in a second trial, with RD = 1 indicating that the cell was active in one trial only and RD = 0 indicating that the firing rates in the two trials were the same.

The RD*_f_* was computed to mitigate the neutralization effect across individual place fields from the same cell:

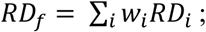

where *RD_i_* is the rate difference of the *i*-th place field and *w_i_* is the weight of the *i*-th place field estimated from its firing rate. RD*_f_* was not computed when changes in firing location occurred.

### Population vector correlation (PVC)

Rate vectors were constructed for the entire population of cells recorded under each condition, including those with firing rates below 0.1 Hz and spatial information content below 0.9586 bits/spike. This process involved arranging all rate maps for all cells in an x-y-z stack, with x and y denoting the two spatial dimensions (20 × 20 bins) and z representing the cell-identity index [41]. For the two-room task, rate maps were rotated analogously to the SC analyses performed for each cell. The distribution of bin rates along the z axis for a specific x-y location represented the composite population vector for that location. Population vectors from corresponding positions were then correlated across pairs of trials. Normalized PVC change was calculated as:

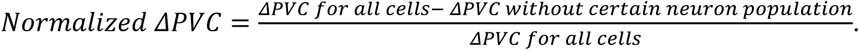

### Shuffling

Shuffling of spatial information was conducted on a cell-by-cell basis, as described previously [37]. Each CA1 cell recorded from the color-reversal task underwent 100 permutations. In each permutation, the entire sequence of spikes fired by the cell was time-shifted along the animal’s path by a random interval ranging from 20 seconds to the total trial length minus 20 seconds. The end of the trial was wrapped back to the beginning to allow for circular displacements. A rate map was generated for each permutation, and the distribution of spatial information values across all permutations for all CA1 cells was determined. A putative excitatory neuron was classified as a place cell if its spatial information content exceeded the 95^th^ percentile of the shuffled distribution (0.9586 bits/spike).

For each CA1 band, a control distribution for SC analyses was established using a random permutation procedure. This procedure involved all active maps (mean firing rates higher than 0.10 Hz) from all cells recorded within that band [43]. For each permutation, two distinct active maps from different cells were randomly selected and correlated in 90° rotation steps to achieve the highest correlation score. In total, 10 000 permutations were performed for each CA1 band. This method preserved the rate maps within each band while eliminating connections between maps. The distribution of random correlations from all 10 000 permutations was then calculated, and the 95^th^ percentile was determined.

### Bayesian decoding

A Bayesian decoding algorithm was employed to convert ensemble spiking activity into positional information for each experiment. The decoding process used a 2 000 ms sliding time window, advancing by 200 ms with each step. The probability of a rat being at position *x*, given the number of spikes *n* from each unit recorded within the time window, was estimated using Bayes’ rule [71]:

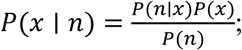

The probabilistic likelihood *P*(*n*|*x*) was calculated using the probability mass function of the Poisson distribution to determine the likelihood of observing a specific number of spikes given the activity pattern of a neuron:

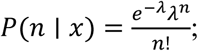

where *λ* is the expected number of spikes based on the tuning curve and *n* is the actual observed number of spikes within the given time window. The prior probability of position *P*(*x*) was set to the frequency of the animal’s presence in each state within the training data. The normalizing constant *P*(*n*) ensured that the posterior probability *P*(*x*|*n*) summed to 1. For visualization and maximum decoding performance estimation, only experiments with over 30 simultaneously recorded place cells in both CA1 bands were considered. Downsampling was applied to equalize cell numbers and repeated three times to determine the mean decoding accuracy value. For position decoding, 90% of the data from the four trials served as the training set, while the remaining 10% was used as the test set. This cross-validation procedure was repeated 10 times (10-fold validation). For context decoding, data from trials 2 and 3 formed the training set, with 90% of trial 1 data serving as the test set. This procedure was also repeated 10 times. Decoding accuracy was expressed as:

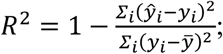

Where *ŷ_i_* and *y_i_* are the predicted and true positions of the *i*-th time bin, respectively, and *ȳ* is the mean position value.

### Fitting

The EzyFit Toolbox in Matlab (v2.44) was used to compute curve fitting for the average data obtained from the tetrodes. The explained variance was calculated as:

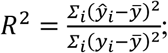

where *ŷ_i_* is the predicted value based on the curve fitting and normalized position of each tetrode, *y_i_* is the average tetrode data, and *ȳ* is the mean value across all tetrodes.

### Statistical procedures

Results are presented as medians and interquartile ranges (IQR) due to the non-normal distribution of most data. The Kruskal-Wallis test was used to compare the firing properties of cells across CA1 bands. The Mann-Whitney U test was employed to compare two groups of unrelated measures, while the Wilcoxon signed-rank test and paired *t*-test were employed to compare two related measures. Regional differences in remapping between two conditions were assessed using two-way repeated-measures ANOVA with a 3 × 2 factor design, considering condition pairs (matching or different) as within-subject factors and CA1 bands as between-subjects factors. Two-way repeated-measures ANOVA was also used when comparing data across recording experiments. The Pearson chi-square test and Fisher’s exact test were applied to compare frequency distributions across groups, and Pearson correlation was calculated to measure the linear correlation between two variables. The binomial test was applied to determine whether the observed frequency differed from the expected frequency. Holm-Bonferroni *post-hoc* tests were used to identify significant differences between pairs. The significance level was set at *P* < 0.05 for two-tailed tests. All statistical analyses were conducted using SPSS v25 (IBM, USA).

### Editing

Kimi AI was used exclusively for language refinement. The authors reviewed and edited the output as needed and take full responsibility for the content of the publication.

## Acknowledgments

The authors express gratitude to Yi-Qing Hui, Yi-Fan Ye, Chang-Lin Shen, and Yi-Ling Shi for their technical assistance in animal care, single-unit isolation and Matlab coding. They also thank Edvard Moser and May-Britt Moser for providing the Behavioral Neurology Toolbox and data from five animals, and Cheng Wang, Xin-Jian Li, Dun Mao, Hai-Bing Xu, and Yong-Gang Yao for their valuable suggestions on the manuscript.

## Author Contributions

LL designed the experiments and analyses; YLD, SYH, XFG, JLL, CD, CJL, RZ, and XC performed the experiments; YLD, SYH, and LL conducted the analyses; YMS carried out the Bayesian decoding. YLD, XC, and LL wrote the manuscript. All authors contributed to discussion and interpretation of the results.

## Code accessibility

All custom-written codes used in the analyses are available upon request.

## Data availability

The data supporting the findings of this study are available from the corresponding author upon reasonable request.

## Supplementary material

Supplementary material is accessible in the online version of this paper.

## References

1. Tulving, E., Episodic memory: from mind to brain. Annu Rev Psychol, 2002. 53: p. 1–25.

2. Sugar, J. and M.B. Moser, Episodic memory: Neuronal codes for what, where, and when. Hippocampus, 2019. 29(12): p. 1190–1205.

3. Colgin, L.L., E.I. Moser, and M.B. Moser, Understanding memory through hippocampal remapping. Trends Neurosci, 2008. 31(9): p. 469–77.

4. Latuske, P., et al., Hippocampal Remapping and Its Entorhinal Origin. Front Behav Neurosci, 2017. 11: p. 253.

5. Rivard, B., et al., Representation of objects in space by two classes of hippocampal pyramidal cells. Journal of General Physiology, 2004. 124(1): p. 9–25.

6. Manns, J.R. and H. Eichenbaum, A cognitive map for object memory in the hippocampus. Learn Mem, 2009. 16(10): p. 616–24.

7. Burke, S.N., et al., The influence of objects on place field expression and size in distal hippocampal CA1. Hippocampus, 2011. 21(7): p. 783–801.

8. Deshmukh, S.S. and J.J. Knierim, Influence of local objects on hippocampal representations: Landmark vectors and memory. Hippocampus, 2013. 23(4): p. 253–67.

9. Vandrey, B., S. Duncan, and J.A. Ainge, Object and object-memory representations across the proximodistal axis of CA1. Hippocampus, 2021. 31(8): p. 881–896.

10. Nagelhus, A., et al., Object-centered population coding in CA1 of the hippocampus. Neuron, 2023. 111(13): p. 2091–2104 e14.

11. Canto, C.B., F.G. Wouterlood, and M.P. Witter, What does the anatomical organization of the entorhinal cortex tell us? Neural Plast, 2008. 2008: p. 381243.

12. Hafting, T., et al., Microstructure of a spatial map in the entorhinal cortex. Nature, 2005. 436(7052): p. 801-806.

13. Ouchi, A. and S. Fujisawa, Predictive grid coding in the medial entorhinal cortex. Science, 2024. 385(67101): p. 776-784.

14. Fyhn, M., et al., Spatial representation in the entorhinal cortex. Science, 2004. 305(5688): p. 1258-64.

15. Sargolini, F., et al., Conjunctive representation of position, direction, and velocity in entorhinal cortex. Science, 2006. 312(5774): p. 758-62.

16. Solstad, T., et al., Representation of geometric borders in the entorhinal cortex. Science, 2008. 322(5909): p. 1865-8.

17. Hoydal, O.A., et al., Object-vector coding in the medial entorhinal cortex. Nature, 2019. 568(7752): p. 400-404.

18. Kropff, E., et al., Speed cells in the medial entorhinal cortex. Nature, 2015. 523(7561): p. 419-24.

19. Tukker, J.J., et al., Microcircuits for spatial coding in the medial entorhinal cortex. Physiol Rev, 2022. 102(2): p. 653–688.

20. Long, J.L. and L. Lu, Dynamic coding in the hippocampus during navigation. Zool Res, 2022. 43(6): p. 1023–1025.

21. Hargreaves, E.L., et al., Major dissociation between medial and lateral entorhinal input to dorsal hippocampus. Science, 2005. 308(5729): p. 1792-4.

22. Huang, X., et al., Distinct spatial maps and multiple object codes in the lateral entorhinal cortex. Neuron, 2023. 111(19): p. 3068–3083 e7.

23. Deshmukh, S.S. and J.J. Knierim, Representation of non-spatial and spatial information in the lateral entorhinal cortex. Front Behav Neurosci, 2011. 5: p. 69.

24. Tsao, A., M.B. Moser, and E.I. Moser, Traces of experience in the lateral entorhinal cortex. Curr Biol, 2013. 23(5): p. 399–405.

25. Igarashi, K.M., et al., Coordination of entorhinal-hippocampal ensemble activity during associative learning. Nature, 2014. 510(7503): p. 143-7.

26. Leitner, F.C., et al., Spatially segregated feedforward and feedback neurons support differential odor processing in the lateral entorhinal cortex. Nat Neurosci, 2016. 19(7): p. 935–44.

27. Tsao, A., et al., Integrating time from experience in the lateral entorhinal cortex. Nature, 2018. 561(7721): p. 57-62.

28. Wang, C., et al., Egocentric coding of external items in the lateral entorhinal cortex. Science, 2018. 362(6417): p. 945-949.

29. Cappaert, N.L., N.M. van Strien, and M.P. Witter, Hippocampal Formation, in The Rat Nervous System, G. Paxinos, Editor. 2015, Academic Press. p. 511-573.

30. Nilssen, E.S., et al., Neurons and networks in the entorhinal cortex: A reappraisal of the lateral and medial entorhinal subdivisions mediating parallel cortical pathways. Hippocampus, 2019. 29(12): p. 1238–1254.

31. Nakamura, N.H., et al., Proximodistal segregation of nonspatial information in CA3: preferential recruitment of a proximal CA3-distal CA1 network in nonspatial recognition memory. J Neurosci, 2013. 33(28): p. 11506–14.

32. Beer, Z., et al., The memory for time and space differentially engages the proximal and distal parts of the hippocampal subfields CA1 and CA3. PLoS Biol, 2018. 16(8): p. e2006100.

33. Flasbeck, V., et al., Spatial information is preferentially processed by the distal part of CA3: Implication for memory retrieval. Behav Brain Res, 2018. 354: p. 31–38.

34. Ito, H.T. and E.M. Schuman, Functional division of hippocampal area CA1 via modulatory gating of entorhinal cortical inputs. Hippocampus, 2012. 22(2): p. 372–87.

35. Nakazawa, Y., et al., Memory retrieval along the proximodistal axis of CA1. Hippocampus, 2016. 26(9): p. 1140–8.

36. Hartzell, A.L., et al., Transcription of the immediate-early gene Arc in CA1 of the hippocampus reveals activity differences along the proximodistal axis that are attenuated by advanced age. J Neurosci, 2013. 33(8): p. 3424–33.

37. Henriksen, E.J., et al., Spatial representation along the proximodistal axis of CA1. Neuron, 2010. 68(1): p. 127–37.

38. Oliva, A., et al., Spatial coding and physiological properties of hippocampal neurons in the Cornu Ammonis subregions. Hippocampus, 2016. 26(12): p. 1593–1607.

39. Deshmukh, S.S., Distal CA1 Maintains a More Coherent Spatial Representation than Proximal CA1 When Local and Global Cues Conflict. J Neurosci, 2021. 41(47): p. 9767–9781.

40. Li, Y., et al., A distinct entorhinal cortex to hippocampal CA1 direct circuit for olfactory associative learning. Nat Neurosci, 2017. 20(4): p. 559–570.

41. Leutgeb, S., et al., Independent codes for spatial and episodic memory in hippocampal neuronal ensembles. Science, 2005. 309(5734): p. 619-23.

42. Lu, L., et al., Impaired hippocampal rate coding after lesions of the lateral entorhinal cortex. Nat Neurosci, 2013. 16(8): p. 1085–93.

43. Lu, L., et al., Topography of place maps along the CA3-to-CA2 axis of the hippocampus. Neuron, 2015. 87(5): p. 1078–92.

44. Burke, S.N., et al., Representation of three-dimensional objects by the rat perirhinal cortex. Hippocampus, 2012. 22(10): p. 2032–44.

45. Deshmukh, S.S., J.L. Johnson, and J.J. Knierim, Perirhinal cortex represents nonspatial, but not spatial, information in rats foraging in the presence of objects: comparison with lateral entorhinal cortex. Hippocampus, 2012. 22(10): p. 2045–58.

46. Aggleton, J.P. and A.J.D. Nelson, Distributed interactive brain circuits for object-in-place memory: A place for time? Brain Neurosci Adv, 2020. 4: p. 2398212820933471.

47. Parron, C. and E. Save, Comparison of the effects of entorhinal and retrosplenial cortical lesions on habituation, reaction to spatial and non-spatial changes during object exploration in the rat. Neurobiology of Learning and Memory, 2004. 82(1): p. 1–11.

48. Rodo, C., F. Sargolini, and E. Save, Processing of spatial and non-spatial information in rats with lesions of the medial and lateral entorhinal cortex: Environmental complexity matters. Behavioural Brain Research, 2017. 320: p. 200–209.

49. Van Cauter, T., et al., Distinct roles of medial and lateral entorhinal cortex in spatial cognition. Cereb Cortex, 2013. 23(2): p. 451–9.

50. Tennant, S.A., et al., Stellate Cells in the Medial Entorhinal Cortex Are Required for Spatial Learning. Cell Reports, 2018. 22(5): p. 1313–1324.

51. Wilson, D.I., et al., Lateral entorhinal cortex is critical for novel object-context recognition. Hippocampus, 2013. 23(5): p. 352–66.

52. Wilson, D.I., et al., Lateral entorhinal cortex is necessary for associative but not nonassociative recognition memory. Hippocampus, 2013. 23(12): p. 1280–90.

53. Ocko, S.A., et al., Emergent elasticity in the neural code for space. Proc Natl Acad Sci U S A, 2018. 115(50): p. E11798–e11806.

54. Wen, J.H., et al., One-shot entorhinal maps enable flexible navigation in novel environments. Nature, 2024.

55. Sun, Y., et al., CA1-projecting subiculum neurons facilitate object-place learning. Nat Neurosci, 2019. 22(11): p. 1857–1870.

56. Sun, Y., et al., Subicular neurons encode concave and convex geometries. Nature, 2024. 627(8005): p. 821-829.

57. Renno-Costa, C., J.E. Lisman, and P.F. Verschure, The mechanism of rate remapping in the dentate gyrus. Neuron, 2010. 68(6): p. 1051–8.

58. Diehl, G.W., et al., Grid and Nongrid Cells in Medial Entorhinal Cortex Represent Spatial Location and Environmental Features with Complementary Coding Schemes. Neuron, 2017. 94(1): p. 83–92 e6.

59. Ismakov, R., et al., Grid Cells Encode Local Positional Information. Curr Biol, 2017. 27(15): p. 2337–2343 e3.

60. Marozzi, E., et al., Purely Translational Realignment in Grid Cell Firing Patterns Following Nonmetric Context Change. Cereb Cortex, 2015. 25(11): p. 4619–27.

61. Miao, C., et al., Hippocampal Remapping after Partial Inactivation of the Medial Entorhinal Cortex. Neuron, 2015. 88(3): p. 590–603.

62. Rueckemann, J.W., et al., Transient optogenetic inactivation of the medial entorhinal cortex biases the active population of hippocampal neurons. Hippocampus, 2016. 26(2): p. 246–60.

63. Kanter, B.R., et al., A Novel Mechanism for the Grid-to-Place Cell Transformation Revealed by Transgenic Depolarization of Medial Entorhinal Cortex Layer II. Neuron, 2017. 93(6): p. 1480–1492 e6.

64. Fyhn, M., et al., Hippocampal remapping and grid realignment in entorhinal cortex. Nature, 2007. 446(7132): p. 190-4.

65. Leutgeb, S., et al., Distinct ensemble codes in hippocampal areas CA3 and CA1. Science, 2004. 305(5688): p. 1295-1298.

66. Knierim, J.J., J.P. Neunuebel, and S.S. Deshmukh, Functional correlates of the lateral and medial entorhinal cortex: objects, path integration and local-global reference frames. Philos Trans R Soc Lond B Biol Sci, 2014. 369(1635): p. 20130369.

67. Soltesz, I. and A. Losonczy, CA1 pyramidal cell diversity enabling parallel information processing in the hippocampus. Nat Neurosci, 2018. 21(4): p. 484–493.

68. Lu, L., et al., Construction of an Improved Multi-Tetrode Hyperdrive for Large-Scale Neural Recording in Behaving Rats. J Vis Exp, 2018(135).

69. Cacucci, F., et al., Theta-modulated place-by-direction cells in the hippocampal formation in the rat. J Neurosci, 2004. 24(38): p. 8265–77.

70. Skaggs, W.E., et al., Theta phase precession in hippocampal neuronal populations and the compression of temporal sequences. Hippocampus, 1996. 6(2): p. 149–72.

71. Glaser, J.I., et al., Machine Learning for Neural Decoding. eNeuro, 2020. 7(4).

