## supplementary figures for "Object-translocation induces event coding in the hippocampus"

### Supporting Information for
### Object-translocation induces event coding in the hippocampus

Ya-Li Duan[1,3,4], Shu-Yi Hu[1,3,4], Xiao-Fan Ge[1,2], Jia-Li Long[1,3], Chu Deng[1,3], Yu-Ming Sun[1,3], Cheng-Ji Li[1,2], Rong Zhang[1,2], Xing Cai[1,2] & Li Lu[1,2,5]

Li Lu


**Fig. S1**

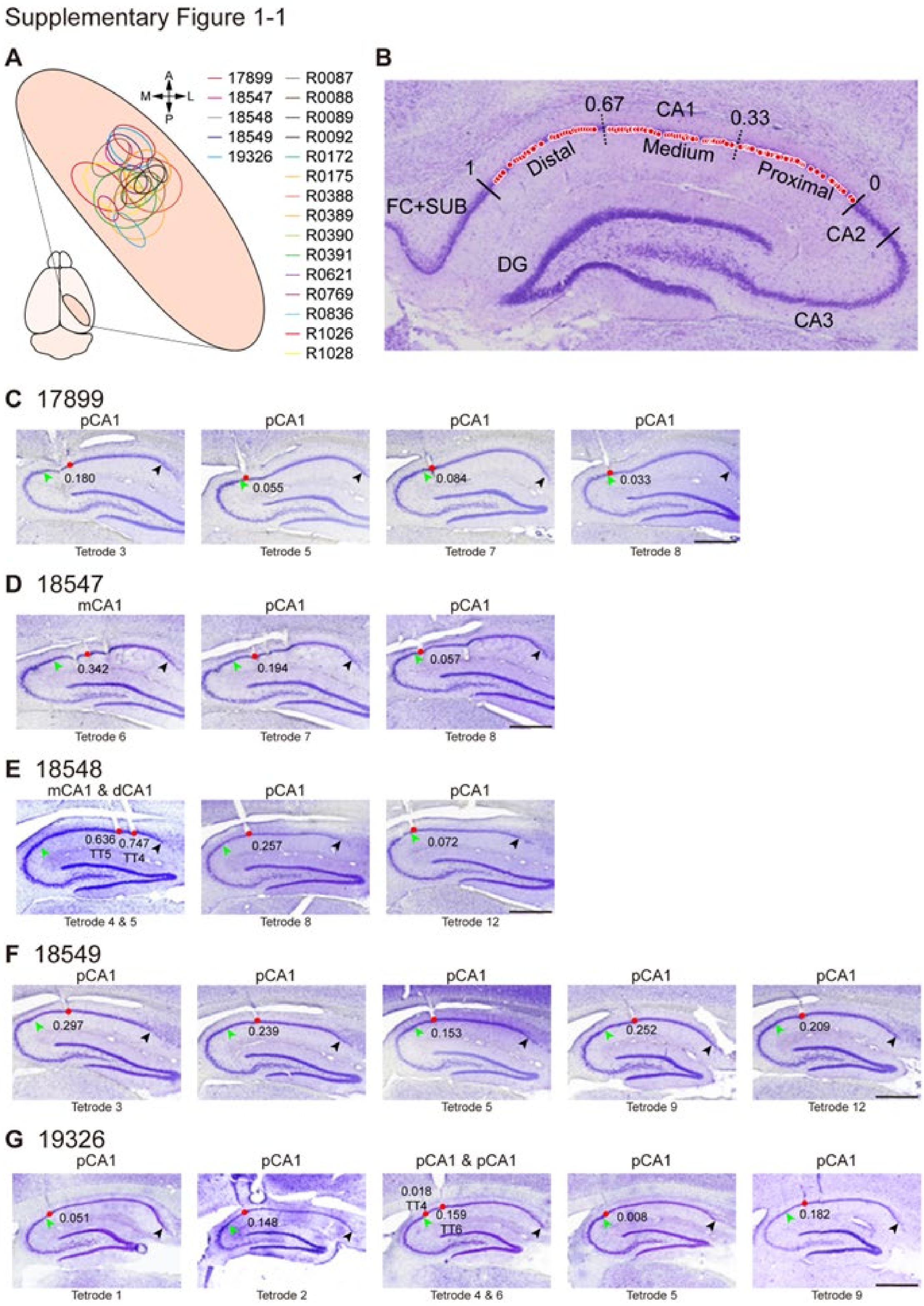
Supplementary Figure 1-1
A
B
17899
18547
18548
18549
19326
R0087
R0088
R0089
R0092
R0172
R0175
R0388
R0389
R0390
R0391
R0621
R0769
R0836
R1026
R1028
0.67
CA1
0.33
1
Distal
Medium
Proximal
0
FC+SUB
CA2
DG
CA3
C 17899
pCA1
0.180
0.055
0.084
0.033
Tetrode 3
Tetrode 5
Tetrode 7
Tetrode 8
D 18547
mCA1
0.342
0.194
0.057
Tetrode 6
Tetrode 7
Tetrode 8
E 18548
mCA1 & dCA1
0.636
TT5
0.747
TT4
0.257
0.072
Tetrode 4 & 5
Tetrode 8
Tetrode 12
F 18549
0.297
0.239
0.153
0.252
0.209
Tetrode 3
Tetrode 5
Tetrode 9
Tetrode 12
G 19326
pCA1 & pCA1
0.051
0.148
0.018
TT4
0.159
TT6
0.008
0.182
Tetrode 1
Tetrode 2
Tetrode 4 & 6
Tetrode 5
Tetrode 9

Supplementary Figure 1-2

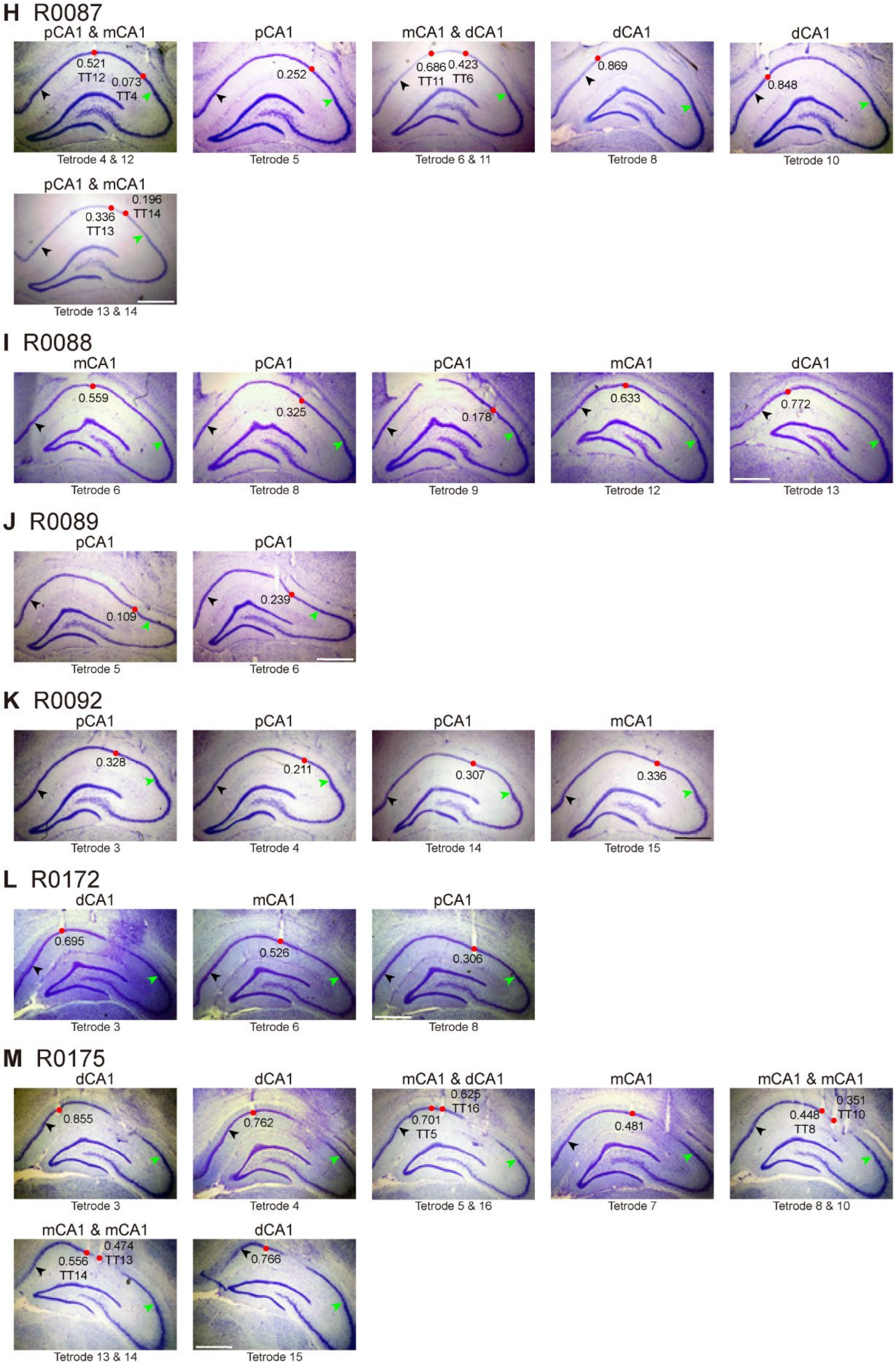

Supplementary Figure 1-3

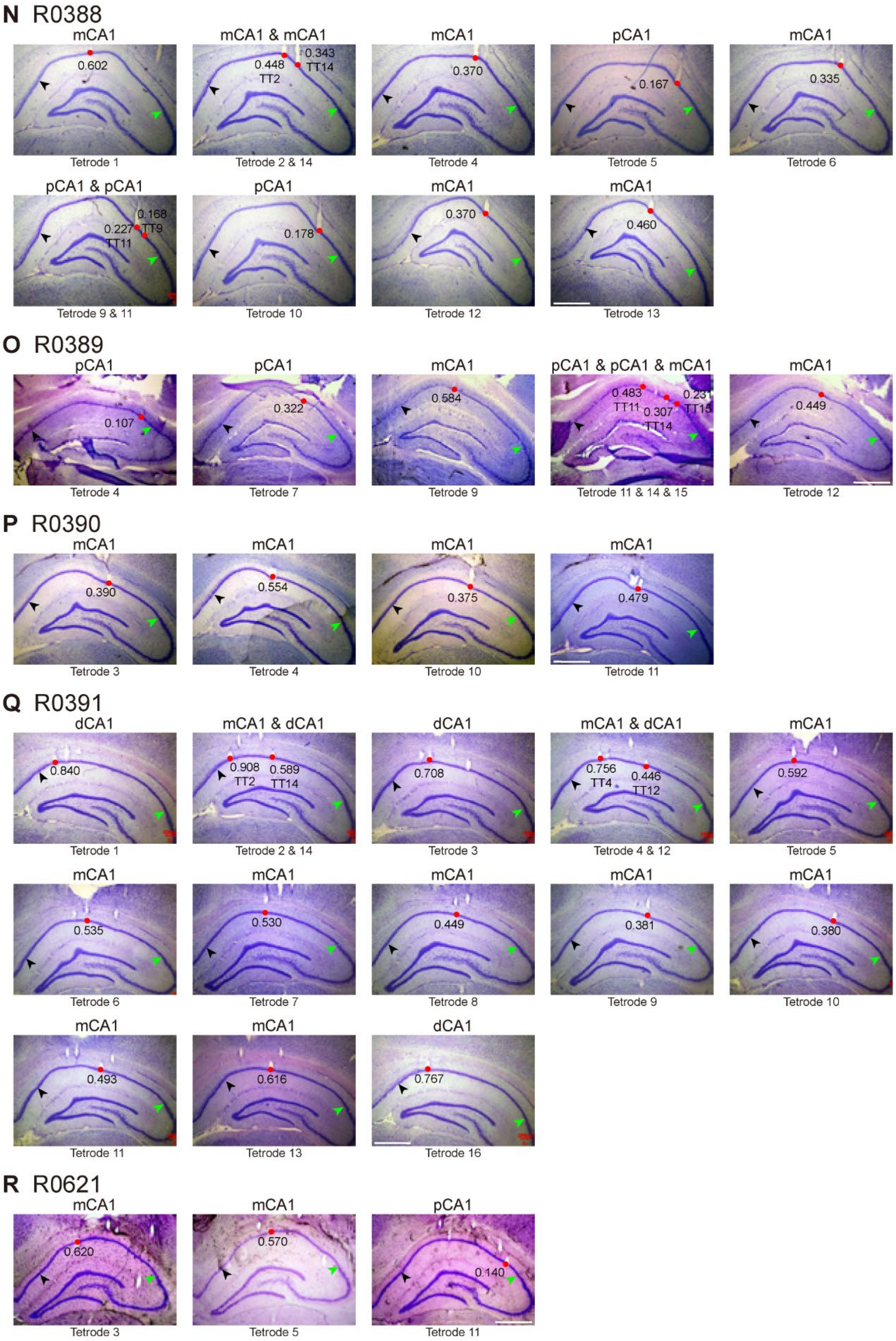

Supplementary Figure 1-4

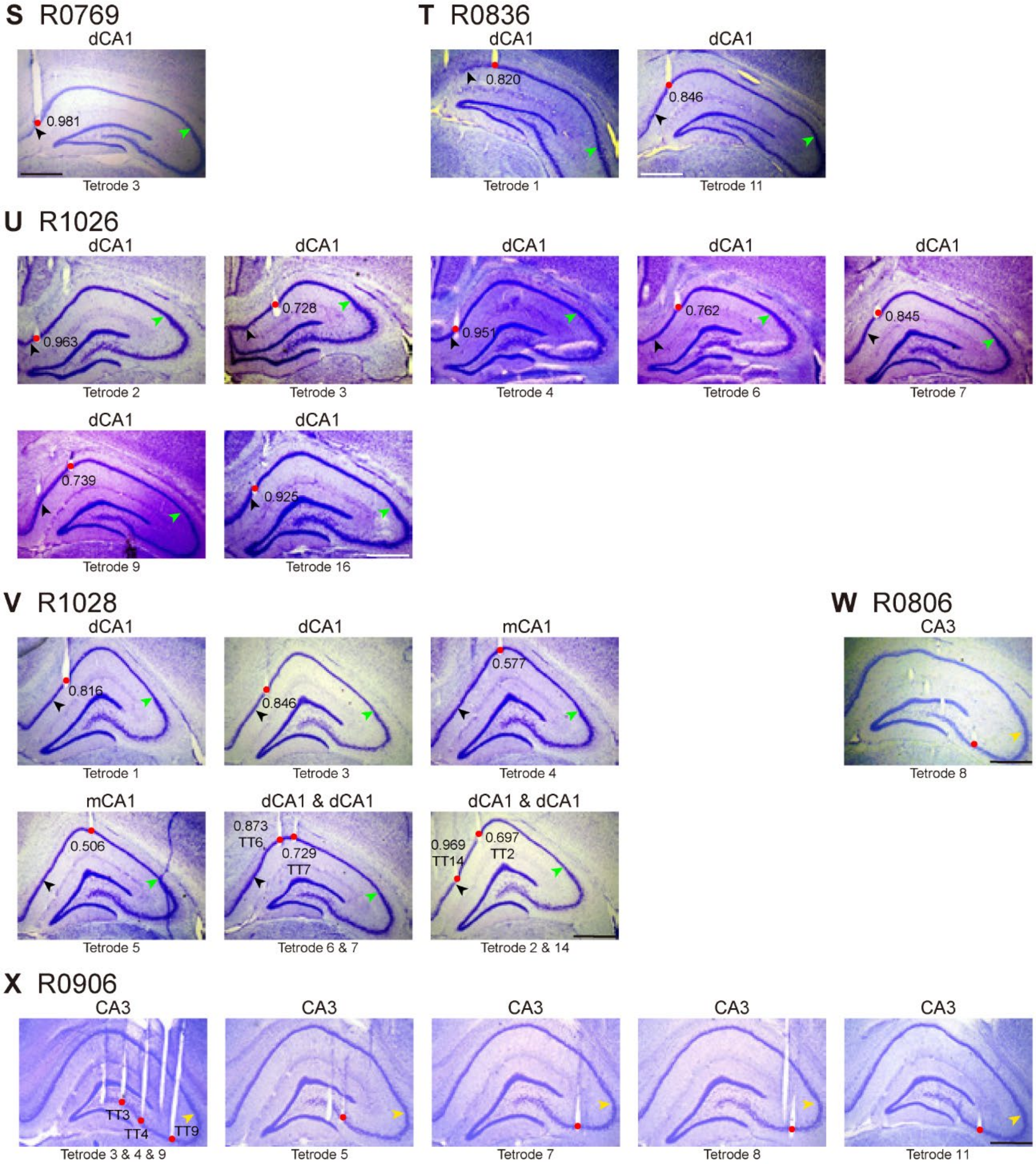


**Figure S1, related to Figure 1. Recording sites in the hippocampus**

**A.** Schematic distribution of estimated recording region in each rat, with color code shown on the right. A, anterior; P, posterior; M: medial; L: lateral.

**B.** Distribution of recording sites along proximodistal axis of CA1. Red dots represent location of tetrode tips. Solid lines indicate borders between hippocampal regions; dashed lines separate proximal, middle, and distal bands of CA1. Numbers indicate normalized positions in CA1 along the transverse axis. FC: fasciola cinereum; SUB: subiculum.

**C–G.** Nissl-stained sagittal sections showing CA1 recording sites from the five animals reported previously [43]. Red dots represent recording sites. Numbers indicate normalized recording positions. Black and green arrowheads indicate the CA1/subiculum and CA2/CA1 borders, respectively. Scale bar: 1 mm.

**H–V.** Nissl-stained coronal sections showing recording sites from the 15 animals recorded in this study. Some tetrodes in R0089 and R0391 were lifted after recording thus recording sites were estimated by the extension of tetrode track. Scale bar: 1 mm.
**W–X.** Nissl-stained coronal sections showing recording sites in CA3 from two rats. Yellow arrowhead indicates the CA3/CA2 border. Scale bar: 1 mm.

**Fig. S2**

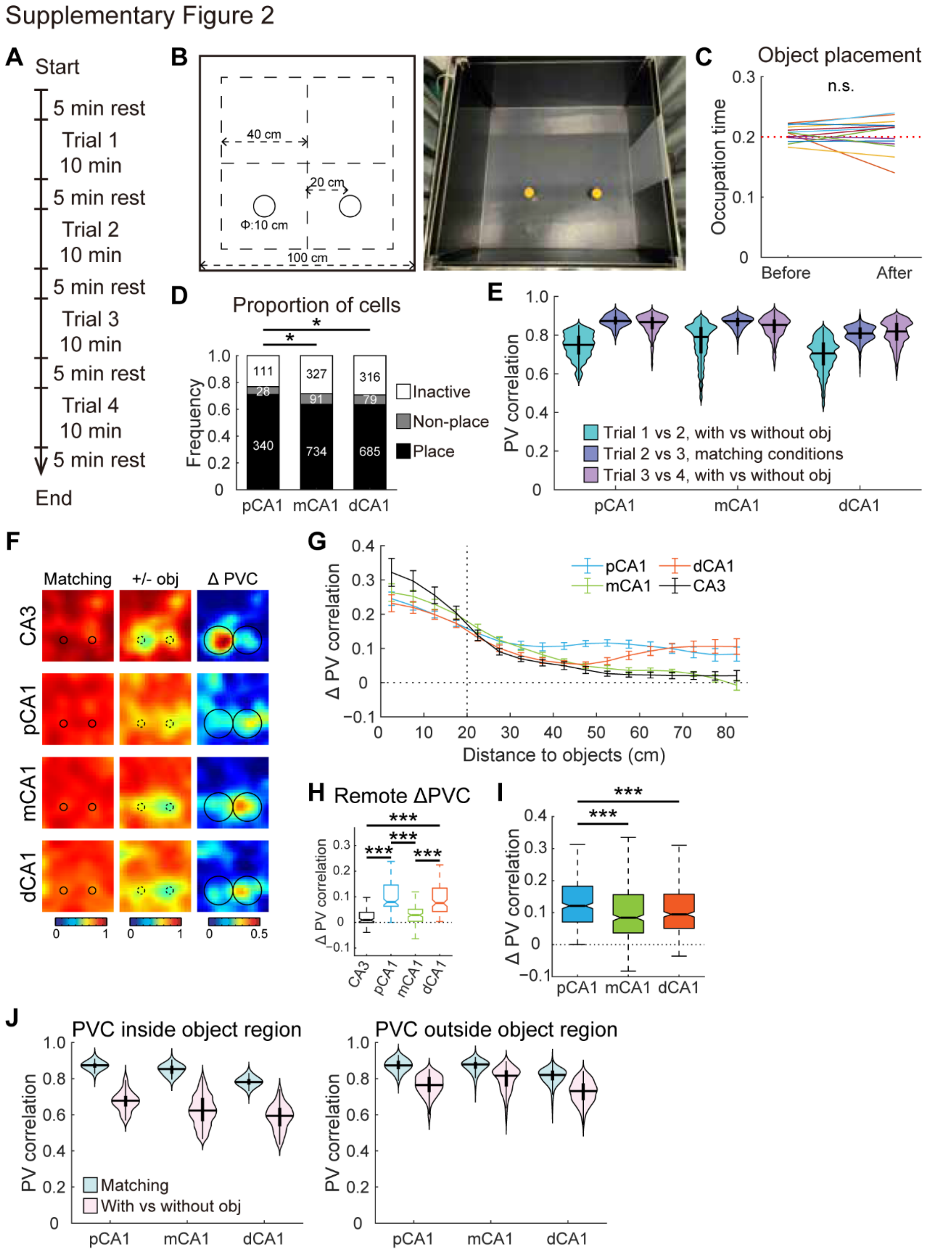


**Figure S2, related to Figure 1. Extended analyses for the object-placement task**

**A.** Timeline of behavioral experiments.

**B.** Objects in the object-placement task.

**C.** Occupation time (OT) in the object region before and after object placement for each rat. The mean OT was close to an unbiased distribution of 0.2 (proportional to region size) and remained unchanged (before: 0.205 ± 0.003; after: 0.204 ± 0.007), paired *t*-test, 14 rats, t = 0.222, $P = 0.828$). n.s., not significant.

**D.** Distribution of pyramidal neurons across three CA1 bands in the task. Inactive cells had firing rates lower than 0.1 Hz. Non-place cells refer to active cells with spatial information scores below the threshold of 0.9586 bits/spike. Numbers indicate cells in each category. Proportion of active cells: pCA1, 76.83%, mCA1, 71.61%, and dCA1: 70.74%; chi-square test, $\chi^2 = 6.449$, $P = 0.040$; Holm-Bonferroni *post hoc* tests, pCA1 vs mCA1, $P = 0.096$; mCA1 vs dCA1, $P = 1.000$; pCA1 vs dCA1, $P = 0.041$. Proportion of place cells: pCA1, 70.98%, mCA1, 63.72%, and dCA1, 63.43%, $\chi^2 = 9.514$, $P = 0.009$; results of *post hoc* tests shown in the plot, *, $P < 0.05$.

**E.** PVCs of CA1 subregions between neighboring trials in the object-placement task. Horizontal line indicates median, and thick vertical bar indicates interquartile range (IQR, 25%–75% percentile) of each subregion. A notable hysteresis effect was observed between trials 3 and 4. Two-way repeated-measures ANOVA for trial pairs 1 vs 2 and 2 vs 3, object: F (1, 1 197) = 2 475.217, $P < 0.001$; band: F (2, 1197) = 182.697, $P < 0.001$); object × band: F (2, 1 197) = 11.762, $P < 0.001$.

**F.** Heatmaps showing PVC of all putative pyramidal neurons in each CA1 subregion, as well as upstream CA3, between trials with or without objects (left two columns), together with changes in PVC between trial pairs (right column). Heatmap bins correspond to spatial bins in rate maps. Color scales are indicated on the right of each row. Small circles represent objects, large circles indicate object region.

**G.** Degree of changes in PVC as a function of distance to the objects (mean ± SEM) in each CA1 subregion, as well as upstream CA3 (301 pyramidal neurons). Both pCA1 and dCA1 showed higher PVC changes distant from the objects.

**H.** Changes in PVC for bins distant (> 50cm) from the objects after object placement, with outliers omitted. CA3: 0.009 (0–0.140), CA2: 0.003 (−0.035–0.089), pCA1: 0.080 (0.063–0.146), mCA1: 0.029 (0.005–0.052), and dCA1: 0.076 (0.043–0.134); Kruskal-Wallis test, 590 bins, H = 189.075, $P < 0.001$. pCA1 and dCA1 were significantly higher than CA3, CA2 and mCA1 (Holm-Bonferroni *post hoc* tests, all $P < 0.001$).

**I.** Degree of changes in PVC between trial pairs, with outliers omitted. Place cells in mCA1 exhibited least activity change. pCA1: 0.121 (0.070–0.183), mCA1: 0.084 (0.037–0.157), and dCA1: 0.095 (0.050–0.158); Kruskal-Wallis test, 1 200 bins, H = 38.042, $P < 0.001$; Holm-Bonferroni *post hoc* tests: ***: $P < 0.001$.

**J.** Correlations of PVs within (left panel) and outside (right panel) the object region, between trials with or without objects. PVC within object region: two-way repeated-measures ANOVA, object: F (1, 237) = 2262.040, $P < 0.001$; band: F (2, 237) = 96.016, $P < 0.001$; object × band: F (2, 237) = 4.720, $P = 0.010$. PVC outside object region: object: F (1, 957) = 2 043.235, $P < 0.001$; band: F (2, 957) = 221.229, $P < 0.001$; object × band: F (2, 957) = 32.592, $P < 0.001$.

Fig. S3

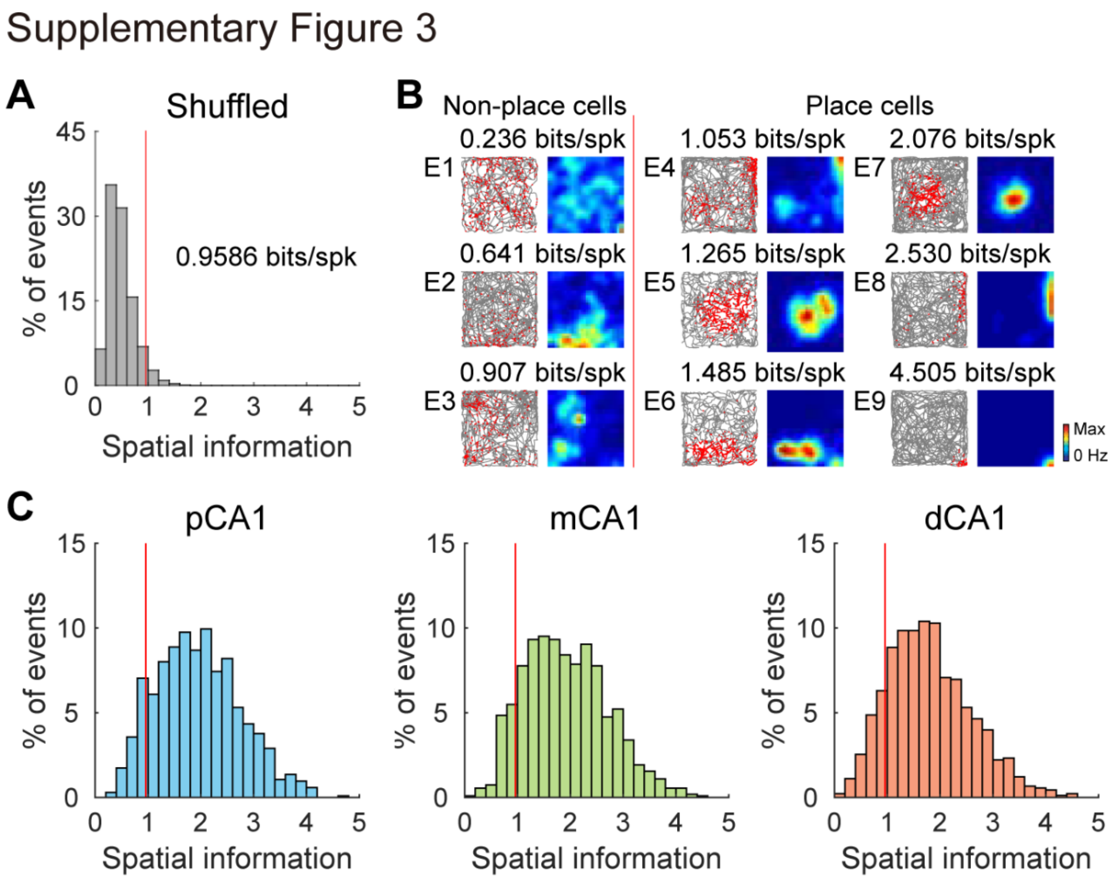


**Figure S3, related to Figure 1. Threshold for place cells**

**A.** Distribution of shuffled spatial information content based on 100 permutations per cell for all CA1 pyramidal neurons. Red line indicates the 95$^{th}$ percentile value (0.9586 bits/spike) for the distribution based on all permutations.

**B.** Example of active cells, aligned according to their spatial information scores. Gray lines indicate trajectories of the animal. Red dots represent individual spikes of each example cell. Color of the rate map is scaled to the peak rate. Spatial information scores are indicated on the top. Red line indicates the threshold for place cells.

**C.** Distribution of spatial information content in pCA1 (left), mCA1 (middle) and dCA1 (right). Red line indicates the threshold for place cells.

**Fig. S4**

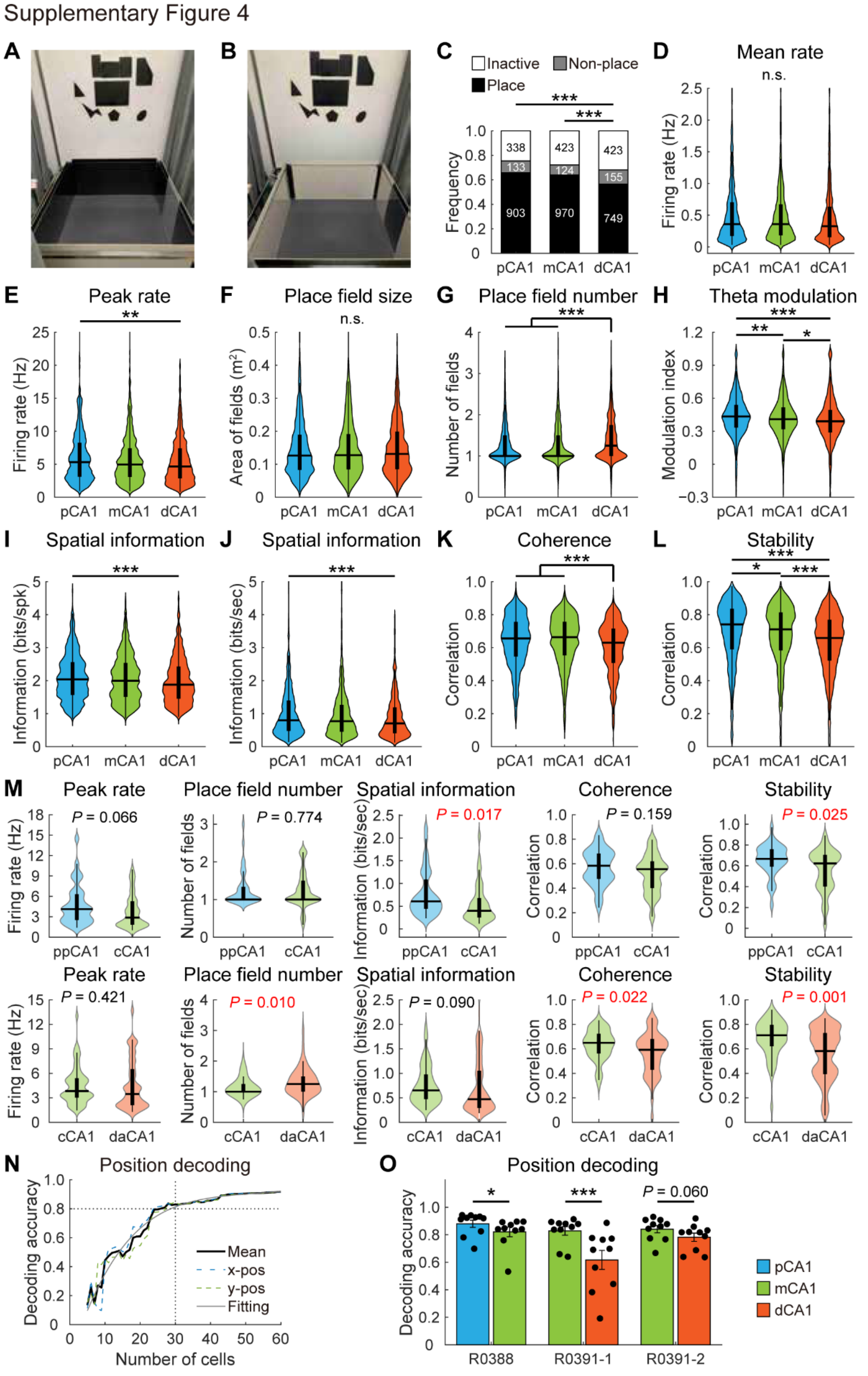
Supplementary Figure 4
A
B
C
Inactive
Non-place
Place
Frequency
pCA1 mCA1 dCA1
D
Mean rate
n.s.
Firing rate (Hz)
E
Peak rate
F
Place field size
Area of fields (m²)
G
Place field number
Number of fields
H
Theta modulation
Modulation index
I
Spatial information
Information (bits/spk)
J
Spatial information
Information (bits/sec)
K
Coherence
Correlation
L
Stability
M
P = 0.066
P = 0.774
P = 0.017
P = 0.159
P = 0.025
ppCA1 cCA1
P = 0.421
P = 0.010
P = 0.090
P = 0.022
P = 0.001
cCA1 daCA1
N
Position decoding
Decoding accuracy
Mean
x-pos
y-pos
Fitting
Number of cells
O
Position decoding
P = 0.060
R0388
R0391-1
R0391-2
pCA1
mCA1
dCA1

**Figure S4, related to Figure 2. Graded spatial tuning along the CA1 transverse axis**

**A.** Test environment with a black recording box and a white cue card. Decorations on the wall provided distal visual cues.

**B.** Test environment with a white recording box and a black cue card.

**C.** Distribution of pyramidal neurons across three CA1 bands in the color-reversal task. Proportion of active cells: pCA1, 75.40%, mCA1, 72.12%, and dCA1: 68.12%; chi-square test, $\chi^2 = 17.744$, $P < 0.001$; Holm-Bonferroni *post hoc* tests, pCA1 vs mCA1, $P = 0.141$; mCA1 vs dCA1, $P = 0.064$; pCA1 vs dCA1, $P < 0.001$. Proportion of place cells: pCA1, 65.72%, mCA1, 63.94%, and dCA1, 56.44%, $\chi^2 = 27.893$, $P < 0.001$; results of *post hoc* tests shown in the plot, ***: $P < 0.001$.

**D**–**L.** Distributions of mean firing rate (**D**), peak firing rate (**E**), size of place field (**F**), number of place field (**G**), theta modulation (**H**), spatial information content (**I**), spatial information rate (**J**), coherence (**K**) and stability (**L**) of place cells recorded in each CA1 band. Mean firing rate (median (25%–75% percentile): pCA1: 0.360 (0.175–0.706) Hz, mCA1: 0.360 (0.183–0.671) Hz, and dCA1: 0.328 (0.155–0.634) Hz; Kruskal-Wallis test, 2 622 cells, H = 3.695, $P = 0.158$. Peak firing rate: pCA1: 5.327 (3.103–8.262) Hz, mCA1: 4.948 (3.065–7.411) Hz, and dCA1: 4.660 (2.854–7.407) Hz; H = 11.566, $P = 0.003$. Place field size: pCA1: 0.126 (0.083–0.189) $m^2$, mCA1: 0.128 (0.084–0.191) $m^2$, and dCA1: 0.131 (0.085–0.199) $m^2$; H = 0.476, $P = 0.788$. Place field number: pCA1: 1.00 (1.00–1.50), mCA1: 1.00 (1.00–1.50), and dCA1: 1.25 (1.00–1.75); H = 28.040, $P < 0.001$. Theta modulation: pCA1: 0.434 (0.333–0.540), mCA1: 0.410 (0.318–0.516), and dCA1: 0.390 (0.290–0.494); H = 28.301, $P < 0.001$. Spatial information content: pCA1: 2.037 (1.565–2.560) bits/spike, mCA1: 1.995 (1.508–2.540) bits/spike, and dCA1: 1.877 (1.454–2.429) bits/spike; H = 13.350, $P = 0.001$. Spatial information rate: pCA1: 0.798 (0.476–1.393) bits/second, mCA1: 0.770 (0.453–1.265) bits/second, dCA1: 0.706 (0.406–1.190) bits/second; H = 14.646, $P = 0.001$. Coherence: pCA1: 0.656 (0.546–0.757), mCA1: 0.664 (0.553–0.758), and dCA1: 0.630 (0.507–0.715); H = 34.163, $P < 0.001$. Stability: pCA1: 0.741 (0.590–0.836), mCA1: 0.711 (0.583–0.814), and dCA1: 0.659 (0.521–0.769); H = 69.309, $P < 0.001$. Holm-Bonferroni *post hoc* tests, n.s.: not significant; **: $P < 0.01$; ***: $P < 0.001$.

**M.** Spatial tuning of place cells recorded from distant proximo-distal locations (normalized distance > 0.1) on the same coronal section. This subset of data compared place cells from disto-anterior (daCA1), central (cCA1), and proximo-posterior portions (ppCA1) of a CA1 transverse block, and exhibited the same trend as the pooled data. Top row: comparison of place cells from ppCA1 and cCA1. Peak firing rate: ppCA1, 4.137 (2.504–6.338) Hz, cCA1, 2.887 (1.839–5.289) Hz; Mann-Whitney U test, 77 cells, Z = 1.838, $P = 0.066$. Place field number: ppCA1, 1.00 (1.00–1.33), cCA1, 1.00 (1.00–1.50); Z = 0.287, $P = 0.774$. Spatial information rate: ppCA1, 0.608 (0.441–1.142) bits/second, cCA1, 0.399 (0.253–0.713) bits/second; Z = 2.389, $P = 0.017$. Coherence: ppCA1, 0.584 (0.476–0.683), cCA1, 0.555 (0.397–0.624); Z = 1.409, $P = 0.159$. Stability: ppCA1, 0.668 (0.589–0.761), cCA1, 0.623 (0.401–0.710); Z = 2.236, $P = 0.025$. Bottom row: comparison of cCA1 and daCA1 place cells. Peak firing rate: cCA1, 3.825 (3.009–5.443) Hz, daCA1, 3.465 (2.101–6.585) Hz; Mann-Whitney U test, 113 cells, Z = 0.805, $P = 0.421$. Place field number: cCA1, 1.00 (1.00–1.25), daCA1, 1.25 (1.00–1.50); Z = −2.567, $P = 0.010$. Spatial information rate: cCA1, 0.652 (0.469–0.978) bits/second, daCA1, 0.470 (0.291–1.069) bits/second; Z = 1.697, $P = 0.090$. Coherence: cCA1, 0.650 (0.562–0.725), daCA1, 0.593 (0.428–0.682); Z = 2.296, $P = 0.022$. Stability: cCA1, 0.711 (0.618–0.798), daCA1, 0.582 (0.394–0.739); Z = 3.366, $P = 0.001$. The observed variations in spatial tuning were not attributed to the dorsoventral distribution of recording sites, as cells recorded from the same transverse block of CA1 displayed a similar trend along the proximodistal axis.

**N.** Decoding accuracy of animal's position as a function of number of cells (n) included in calculation. Data were generated by taking a random subset of n cells from a total population of over 60 simultaneously recorded place cells. The mean accuracy values averaged from 10-fold crossvalidations for the 60 loops are represented. The decoding accuracy reached 0.8 when 30 cells were included.

**O.** Position decoding accuracy (mean ± SEM) for simultaneously recorded pCA1/mCA1 and mCA1/dCA1 data sets. Black dots represent data from individual predictions. The three experiments with more than 30 simultaneously recorded place cells in both bands are shown. Downsampling was applied to equalize cell numbers. R0388: pCA1: 0.880 ± 0.026; mCA1: 0.821 ± 0.035; paired *t*-test, t = 3.131, $P$ = 0.012, 10 calculations. R0391-1: mCA1: 0.828 ± 0.031; dCA1: 0.617 ± 0.069; t = 4.587, $P$ = 0.001. R0391-2: mCA1: 0.841 ± 0.026; dCA1: 0.782 ± 0.030; t = 2.149, $P$ = 0.060. *: $P < 0.05$; ***: $P \leq 0.001$.

**Fig. S5**

Supplementary Figure 5

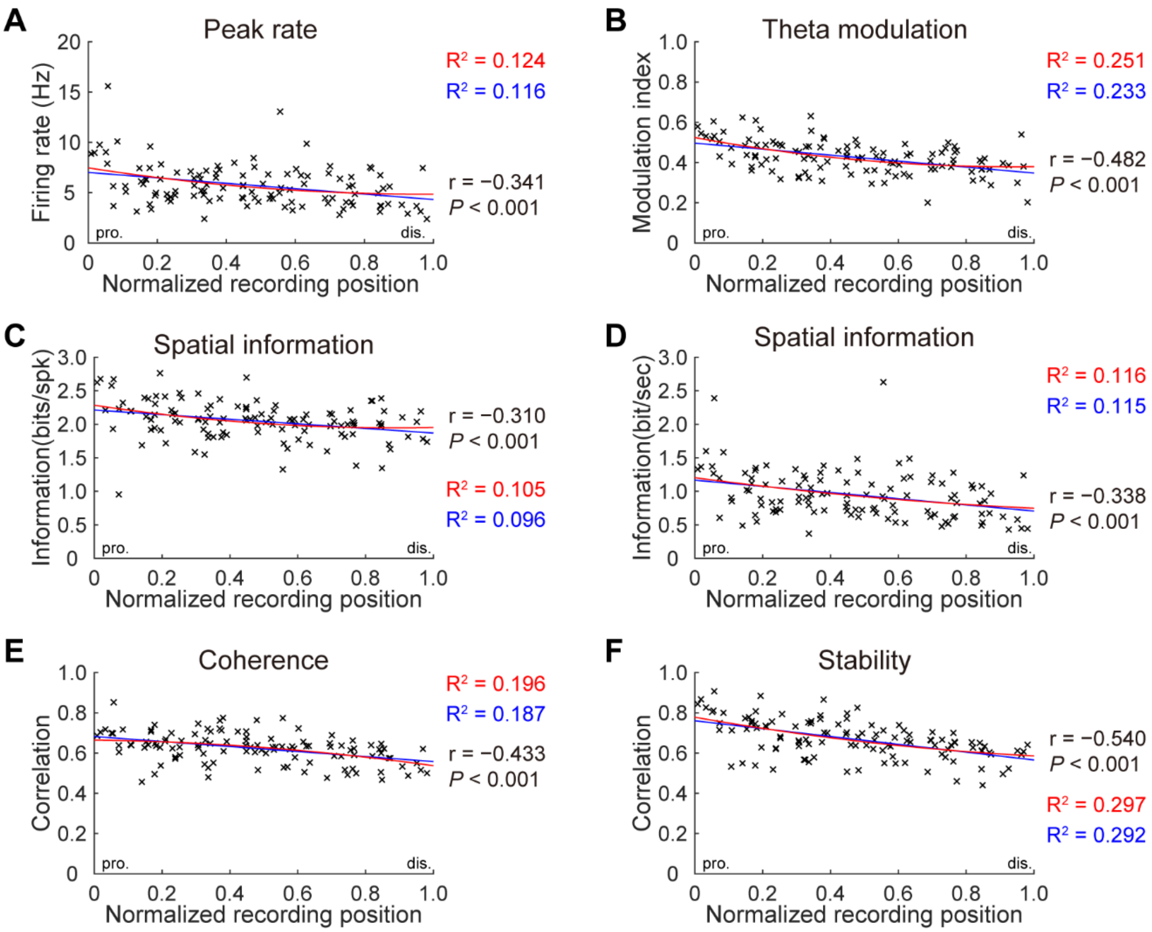


**Figure S5, related to Figure 2. Linear reduction in spatial tuning from pCA1 to dCA1**

Correlations between normalized recording position (113 tetrodes) and peak rate (**A**), theta modulation (**B**), spatial information content (**C**), spatial information rate (**D**), coherence (**E**), and stability (**F**). Each cross corresponds to average data of a single tetrode. Blue curve, linear regression; red curve, quadratic polynomial regression. Explained variances ($R^2$) are indicated with the same color code, with Pearson correlation coefficients and *P* values provided. pro., proximal end; dis., distal end.

**Fig. S6**

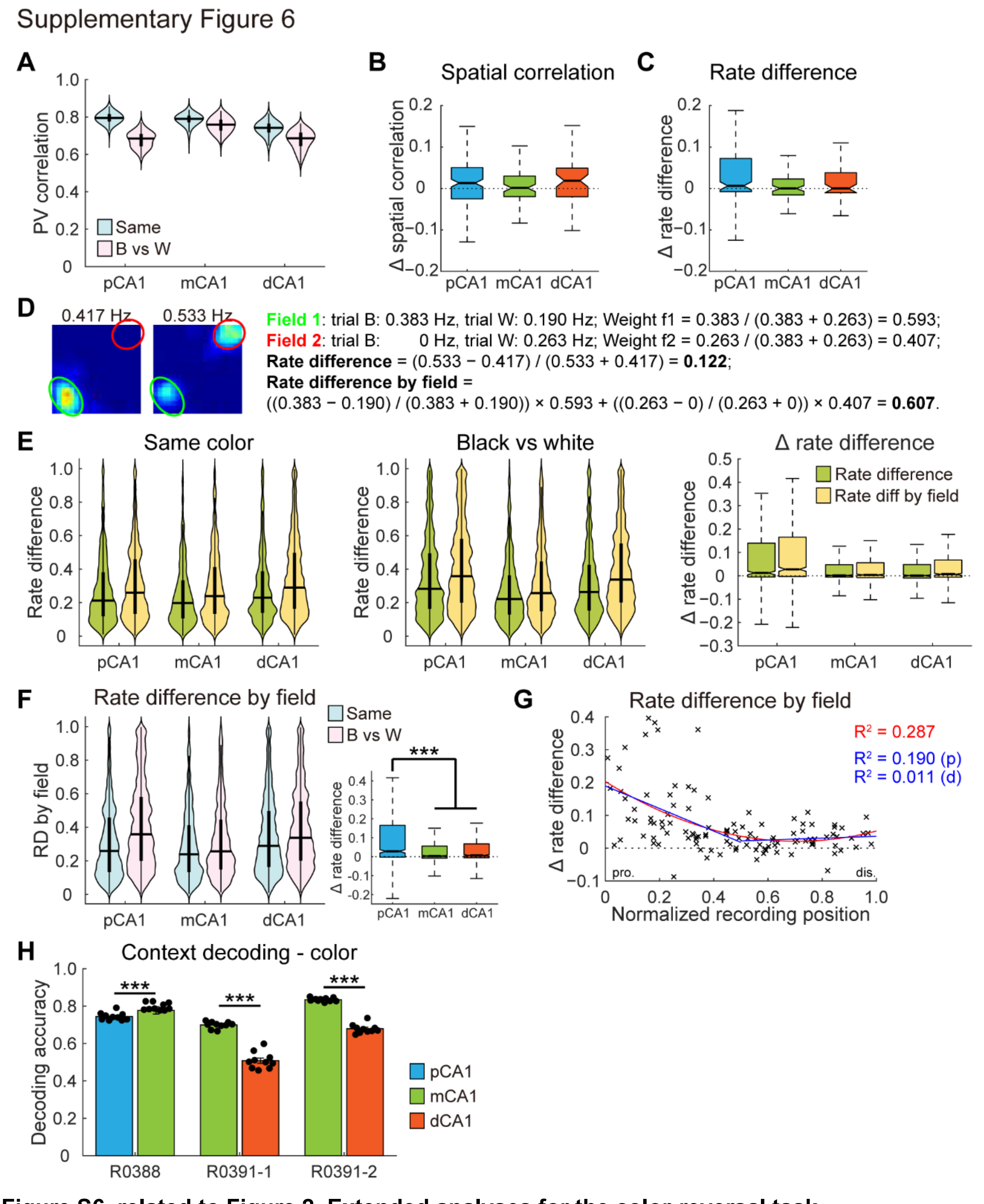


**Figure S6, related to Figure 2. Extended analyses for the color-reversal task**

**A.** Distribution of PVC in the same color condition (same) and the color-reversed condition (B vs W) across CA1 subregions. Two-way repeated-measures ANOVA, color: F (1, 1 197) = 3 336.027, $P < 0.001$; band: F (2, 1 197) = 396.174, $P < 0.001$; color × band: F (2, 1 197) = 448.356, $P < 0.001$. **B**–**C.** Similar V-shaped pattern was observed in SC (**B**) and RD (**C**) when only simultaneous recordings from all three CA1 subregions were included for analysis. SC: pCA1: 0.013 (−0.025–0.051), mCA1: 0 (−0.016–0.023), and dCA1: 0.019 (−0.020–0.049); Kruskal-Wallis test, 291 cells, H = 1.513, $P = 0.469$. RD: pCA1: 0.006 (−0.008–0.072), mCA1: 0.001 (−0.020–0.030), and dCA1:

0 (−0.010–0.038); H = 6.687, $P$ = 0.035.

**D.** Representative rate maps from a place cell with two individually remapped firing fields in black (top) and white (bottom) conditions. Numbers on top indicate overall firing rate in the trial. The two fields remapped independently in reverse directions so that the changes in firing rate were neutralized with the measure of RD (RD) but not with RD by field ($RD_f$).

**E.** Comparison of RD and $RD_f$ across CA1 subregions. Although $RD_f$ were significantly higher than RD between conditions (Same: two-way repeated measures ANOVA, measure: F (1, 2 619) = 388.282, $P$ < 0.001; band: F (2, 2 619) = 14.263, $P$ < 0.001; measure × band: F (2, 2 619) = 6.952, $P$ = 0.001; B vs W: measure: F (1, 2 619) = 533.091, $P$ < 0.001; band: F (2, 2 619) = 37.929, $P$ < 0.001; measure × band: F (2, 2 619) = 5.481, $P$ = 0.004), elevations in $RD_f$ between black and white conditions were marginal (measure: F (1, 2 619) = 36.765, $P$ < 0.001; band: F (2, 2 619) = 63.833, $P$ < 0.001; measure × band: F (2, 2 619) = 11.101, $P$ < 0.001).

**F.** Left: distributions of $RD_f$ in CA1 subregions between matching (Same) and color reversed (B vs W) conditions in the color-reversal task. Horizontal line indicates median, and thick vertical bar indicates IQR of each subregion. A pattern similar to change in overall RD was observed with the measure of $RD_f$ (two-way repeated measures ANOVA, color: F (1, 2 619) = 397.067, P < 0.001; band: F (2, 2 619) = 23.230, P < 0.001; color × band: F (2, 2 619) = 67.922, P < 0.001). Right: changes in $RD_f$ between condition pairs, with outliers omitted pCA1: 0.028 (−0.003–0.166), mCA1: 0.003 (−0.008–0.056), and dCA1: 0.006 (−0.006–0.067); Kruskal-Wallis test, 2 622 cells, H = 65.433, $P$ < 0.001, Holm-Bonferroni *post hoc* tests, ***: $P$ < 0.001.

**G.** Changes in $RD_f$ between condition pairs as a function of tetrode position. Each cross corresponds to average data of a single tetrode. Blue curve, linear regression; red curve, quadratic polynomial regression. Explained variances ($R^2$) are indicated with the same color code. pro., proximal end; dis., distal end. p, proximal half; d, distal half. Change in $RD_f$ was significantly correlated with tetrode location in the proximal half of CA1, but not in the distal half (Pearson correlation: proximal: r = −0.435, $P$ < 0.001; distal: r = 0.106, $P$ = 0.462).

**H.** Context decoding accuracy (mean ± SEM) for the same data sets from Figure S4O. Black dots represent data from individual predictions. R0388: pCA1, 0.744 ± 0.007, mCA1, 0.797 ± 0.006; paired *t*-test, t = 6.714, $P$ < 0.001, 10 calculations. R0391-1: mCA1, 0.699 ± 0.006, dCA1, 0.508 ± 0.014; t = 14.98, $P$ < 0.001. R0391-2: mCA1, 0.834 ± 0.003, dCA1, 0.678 ± 0.008); t = 27.95, $P$ < 0.001. ***: $P$ < 0.001.

**Fig. S7**

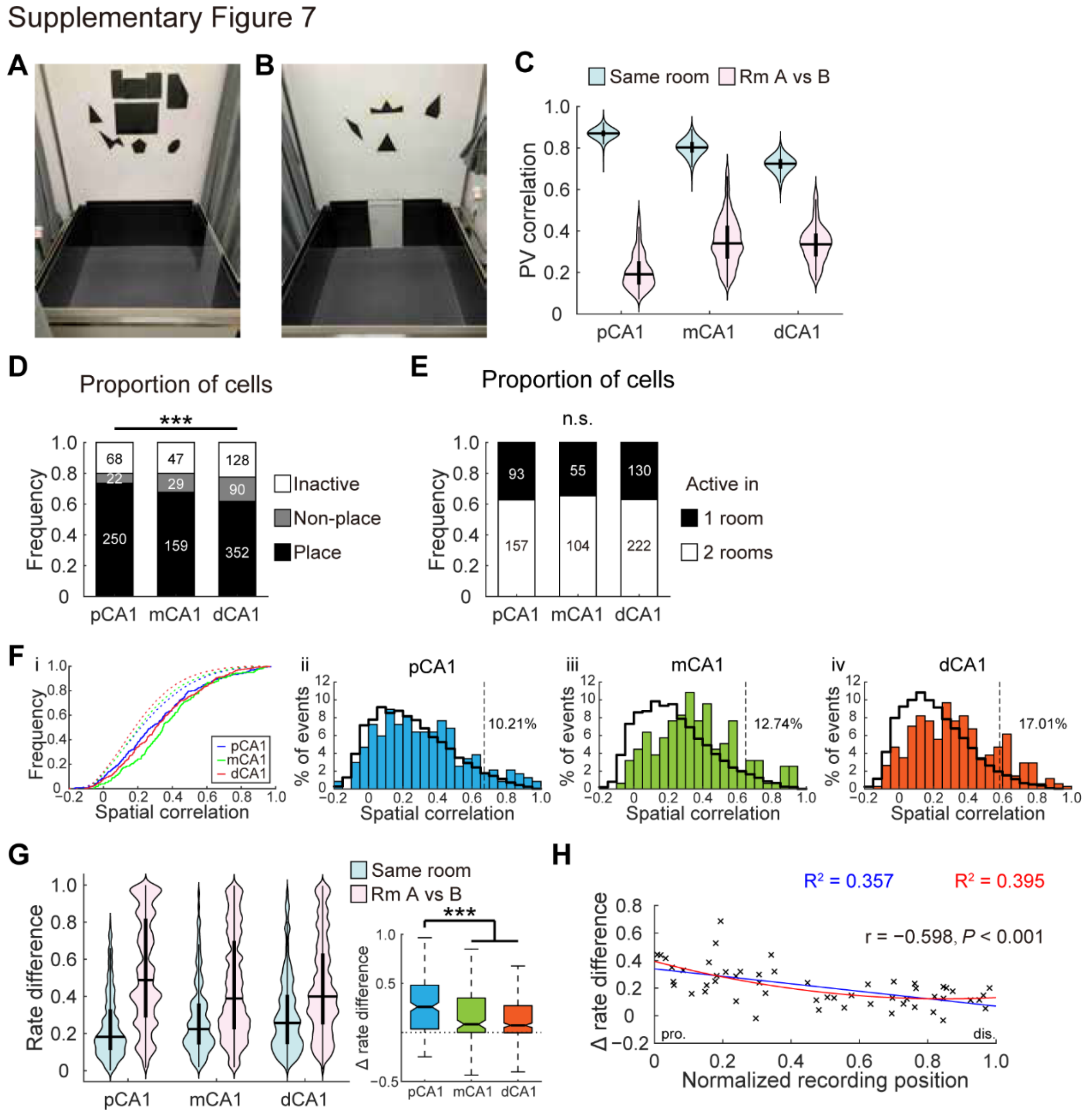


**Figure S7, related to Figure 3. Extended analyses for the two-room task**

**A–B.** Test environments in room A (**A**) and B (**B**), with distinct decoration patterns on the room wall. To further discriminate the two recording rooms, the cue card in room B was positioned 90 degrees counterclockwise relative to room A.

**C.** Distribution of PVC in CA1 subdivisions in the same room (Same) and in different rooms (Room A vs B) in the two-room task. Two-way repeated-measures ANOVA, room: F (1, 1 197) = 28 489.675, $P < 0.001$; band: F (2, 1 197) = 83.437, $P < 0.001$; room × band: F (2, 1 197) = 800.992, $P < 0.001$.

**D.** Distribution of pyramidal neurons across three CA1 bands, from rats exhibiting global remapping in the two-room task. Proportion of active cells: pCA1, 80.00%, mCA1, 80.00%, and dCA1: 77.54%; chi-square test, $\chi^2 = 1.033$, $P = 0.598$. Proportion of place cells: pCA1, 73.53%, mCA1, 67.66%, and dCA1, 61.75%, $\chi^2 = 13.437$, $P = 0.001$; results of Holm-Bonferroni *post hoc* tests shown in the plot, ***, $P < 0.001$.

**E.** Distribution of active place cells in the two-room task, with cell numbers indicated. Proportion of cells that were active in both rooms: pCA1, 62.80%; mCA1, 65.41%; and dCA1, 63.07%, chi-square test, $\chi^2 = 0.331$, $P = 0.847$.

**F.** i. Distributions of SC between rate maps for the same cell in different rooms (solid curve) and randomly selected rate maps from different cells (10 000 permutations, dashed curve). ii–iv. Same distributions as in (i), but for individual CA1 subregions. Colored bars show the distribution of SC between rooms; black trace represents the distribution of permutations. Dashed vertical line indicates the 95th percentile value of the latter distribution (chance level). Inter-room correlations tended to skew toward higher values than expected from the chance distribution from pCA1 to dCA1. The proportion of observations above the chance level is indicated on the right. pCA1, 10.21%, mCA1, 12.74%, and dCA1, 17.01%, all significantly higher than chance (5%, all $P \leq 0.001$, binomial tests). There is a considerable trend toward significance from pCA1 to dCA1, chi-square test, $\chi^2 = 5.595$, $P = 0.063$.

**G.** Left: distribution of RD in CA1 subdivisions in the same room, and in different rooms (Room A vs B). Two-way repeated-measures ANOVA, room: $F(1, 758) = 410.559$, $P < 0.001$; band: $F(2, 758) = 0.380$, $P = 0.684$; room × band: $F(2, 758) = 23.136$, $P < 0.001$), Right: change in RD between condition pairs, with outliers omitted. pCA1: 0.261 (0.034–0.484), mCA1: 0.082 (0–0.352), and dCA1: 0.073 (−0.003–0.274); Kruskal-Wallis test, 761 cells, H = 39.143, $P < 0.001$; Holm-Bonferroni *post hoc* tests: ***: $P < 0.001$.

**H.** Correlation between tetrode position and changes in RD across condition pairs (54 tetrodes). Each cross corresponds to average data of a single tetrode. Blue curve, linear regression; red curve, quadratic polynomial regression.

**Fig. S8**

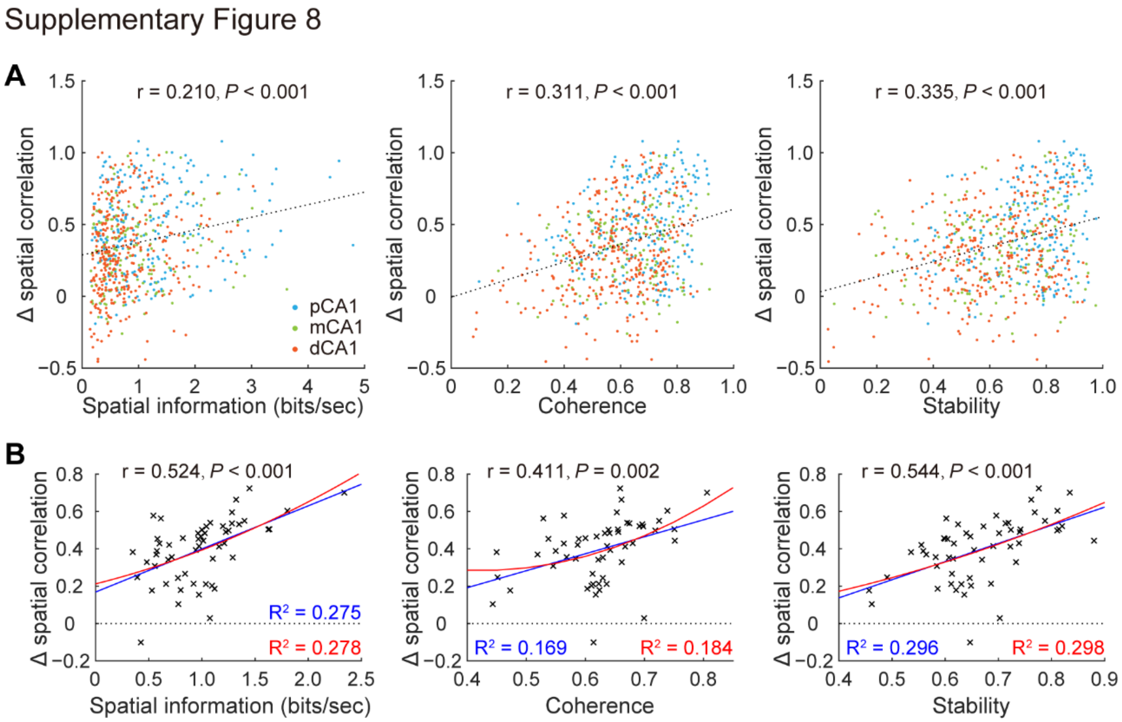


**Figure S8, related to Figure 3. Degree of spatial tuning and sensitivity to prominent landmark changes were correlated**

**A**. Scatter plot showing Pearson correlation between spatial information rate (left), coherence (middle), and stability (right) of individual neurons and changes in SC in the two-room task. Each dot represents data from an individual place cell. Neurons located in pCA1, mCA1 and dCA1 are color coded. Dashed line: linear regression.

**B**. Scatter plot showing Pearson correlation between spatial information rate (left), coherence (middle), and stability (right) and changes in SC in the two-room task, based on data from 54 tetrodes. Each cross corresponds to average data of a single tetrode. Blue curve, linear regression; red curve, quadratic polynomial regression.

**Fig. S9**

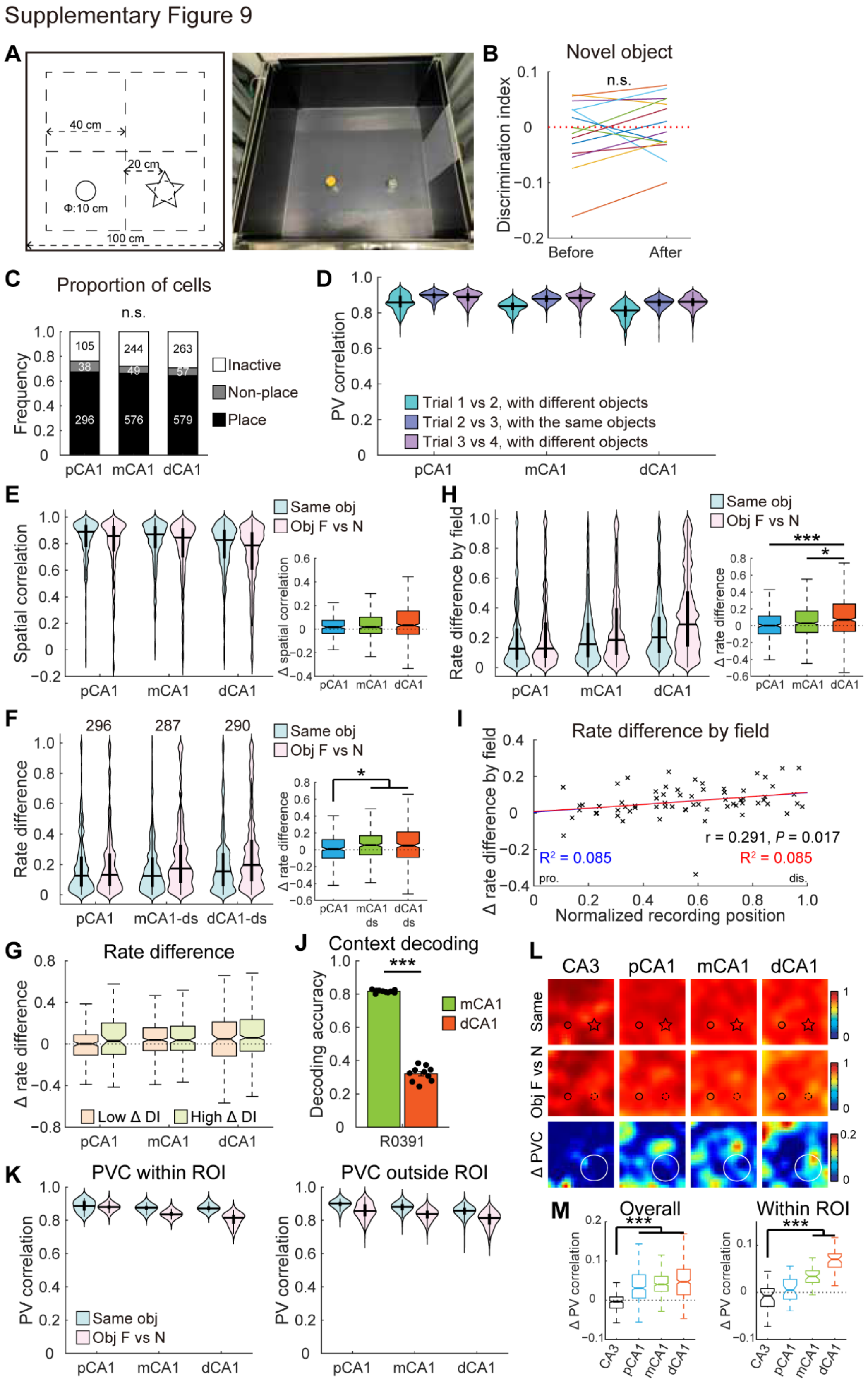

Supplementary Figure 9
A
40 cm
20 cm
Φ:10 cm
100 cm
B
Novel object
n.s.
Discrimination index
Before
After
C
Proportion of cells
n.s.
Frequency
Inactive
Non-place
Place
pCA1
mCA1
dCA1
D
PV correlation
Trial 1 vs 2, with different objects
Trial 2 vs 3, with the same objects
Trial 3 vs 4, with different objects
E
Spatial correlation
Same obj
Obj F vs N
Δ spatial correlation
H
Rate difference by field
Δ rate difference
F
Rate difference
mCA1-ds
dCA1-ds
I
Rate difference by field
Δ rate difference by field
r = 0.291, P = 0.017
R² = 0.085
Normalized recording position
G
Rate difference
Δ rate difference
Low Δ DI
High Δ DI
J
Context decoding
Decoding accuracy
R0391
L
CA3
Same
Obj F vs N
Δ PVC
K
PVC within ROI
PVC outside ROI
M
Overall
Within ROI
Δ PV correlation

**Figure S9, related to Figure 4. Extended analyses for the novel-object task**

**A.** Object replacement in the novel-object task.

**B.** Discrimination index (DI) between the two objects before and after object replacement for each rat. The mean DI was close to unbiased exploration and remained unchanged (before: −0.011 ± 0.016; after: 0.004 ± 0.014, paired *t*-test, 14 rats, t = −1.203, $P = 0.250$). n.s., not significant.

**C.** Distribution of pyramidal neurons across three CA1 bands in the task. Proportion of active cells: pCA1, 76.08%, mCA1, 71.92%, and dCA1: 70.75%; chi-square test, $\chi^2 = 4.279$, $P = 0.117$. Proportion of place cells: pCA1, 67.43%, mCA1, 66.28%, and dCA1, 64.40%, $\chi^2 = 1.379$, $P = 0.505$. n.s., not significant.

**D.** Distribution of overall PVCs of CA1 subregions between neighboring trials in the novel-object task. A notable hysteresis effect was observed between trials 3 and 4. Two-way repeated-measures ANOVA for trial pairs with familiar objects (Trials 2 vs 3), and with different objects (Trials 1 vs 2). novelty: F (1, 1 197) = 1 388.100, $P < 0.001$; band: F (2, 1 197) = 261.805, $P < 0.001$; novelty × band: F (2, 1 197) = 12.330, $P < 0.001$.

**E.** Left: distributions of SC in CA1 subregions between trials with a novel object (Same obj) and trials before and after object replacement (obj F vs N). Two-way repeated-measures ANOVA, novelty: F (1, 1 448) = 69.447, $P < 0.001$; band: F (2, 1 448) = 21.964, $P < 0.001$; novelty × band: F (2, 1 448) = 0.528, $P = 0.590$. Right: changes in activity measures between condition pairs, with outliers omitted. pCA1, 0.016 (−0.034–0.076), mCA1, 0.017 (−0.034–0.101), and dCA1, 0.031 (−0.043–0.154); Kruskal-Wallis test, H = 3.378, $P = 0.151$, 1 451 cells.

**F.** Left: distribution of downsampled RD in CA1 subregions between trial pairs in the task. Numbers on top indicate cell number after downsampling. Similar results were obtained with downsampled data. Two-way repeated measures ANOVA, novelty: F (1, 871) = 42.556, $P < 0.001$; band: F (2, 871) = 4.742, $P = 0.009$; novelty × band: F (2, 871) = 4.637, $P = 0.010$. Right: changes in activity measures between condition pairs based on downsampled data, with outliers omitted. RD: pCA1, 0.008 (−0.101–0.120), mCA1, 0.056 (−0.058–0.166), and dCA1, 0.050 (−0.0890.212); Kruskal-Wallis test, 874 cells, H = 10.954, $P = 0.004$. Holm-Bonferroni *post hoc* tests, *: $P < 0.05$.

**G.** Changes in RD between trial pairs, analyzed separately according to DI change in the object region, with outliers omitted. Low Δ DI: below median of DI change; High Δ DI: above median of DI change. No difference was observed in each CA1 subregion. Mann-Whitney U tests, all $P > 0.05$.

**H.** Left: distribution of $RD_f$ across CA1 subregions between trials with a novel object (Same obj) and trials before and after object replacement (obj F vs N) in the task. Two-way repeated measures ANOVA, novelty: F (1, 1 448) = 63.640, $P < 0.001$; band: F (2, 1 448) = 28.451, $P < 0.001$; novelty × band: F (2, 1 448) = 6.456, $P = 0.002$. Right: degree of changes in $RD_f$ between condition pairs, with outliers omitted. pCA1: 0.002 (−0.095–0.115), mCA1: 0.029 (−0.080–0.175), and dCA1: 0.072 (−0.068–0.260); Kruskal-Wallis test, 1 451 cells, H = 18.747, $P < 0.001$. Holm-Bonferroni *post hoc* tests, *: $P < 0.05$; ***: $P < 0.001$.

**I.** Correlation between normalized recording position and changes in $RD_f$ between condition pairs. Each cross corresponds to average data of a single tetrode. Blue curve, linear regression; red curve, quadratic polynomial regression.

**J.** Context decoding accuracy (mean ± SEM) for the data set with simultaneously recorded mCA1/dCA1 neurons. Black dots represent data from individual predictions. Only one experiment had more than 30 simultaneously recorded place cells in both CA1 subregions. Downsampling was applied to equalize cell numbers. R0391: mCA1, 0.816 ± 0.003, dCA1, 0.321 ± 0.014; paired *t*-test, t = 38.89, $P < 0.001$, 10 calculations. ***: $P < 0.001$.

**K.** Correlations of PVs within (left panel) and outside (right panel) the target region, between trials with familiar objects and trials before and after object replacement. PVC within object region: two-way repeated measures ANOVA, novelty: F (1, 117) = 236.315, $P < 0.001$; band: F (2, 117) = 39.954,

*P* < 0.001; novelty × band: F (2, 117) = 58.668, *P* < 0.001. PVC outside object region: novelty: F (1, 1 077) = 1253.175, *P* < 0.001; band: F (2, 1 077) = 231.473, *P* < 0.001; novelty × band: F (2, 1 077) = 4.168, *P* = 0.016.

**L.** Heatmaps showing PVC of all putative pyramidal neurons in CA3 (293 neurons) between trials (left two columns), together with changes in PVC between trial pairs (right column) in the novel-object task. Small circle and star represent familiar and novel objects, respectively; large circle indicates region of the novel object.

**M.** Degree of changes in overall PVC and PVC with in ROI between trial pairs in CA3 and CA1 subregions, with outliers omitted. Overall change: CA3, −0.003 (−0.020–0.009), pCA1, 0.031 (0.006–0.065), mCA1, 0.040 (0.023–0.061), and dCA1, 0.047 (0.014–0.079). Kruskal-Wallis test, 1600 bins, H = 429.623, P < 0.001. CA3 was significantly lower than all CA1 subregions (Holm-Bonferroni *post hoc* tests, all *P* < 0.001). Changes within ROI: CA3, −0.007 (−0.030–0.008), pCA1, 0.005 (−0.014–0.028), mCA1, 0.034 (0.021–0.046), and dCA1, 0.070 (0.053–0.082). H = 90.996, P < 0.001, 160 bins. CA3 was significantly lower than mCA1 and dCA1 (*post hoc* tests, both *P* < 0.001).

**Fig. S10**

Supplementary Figure 10

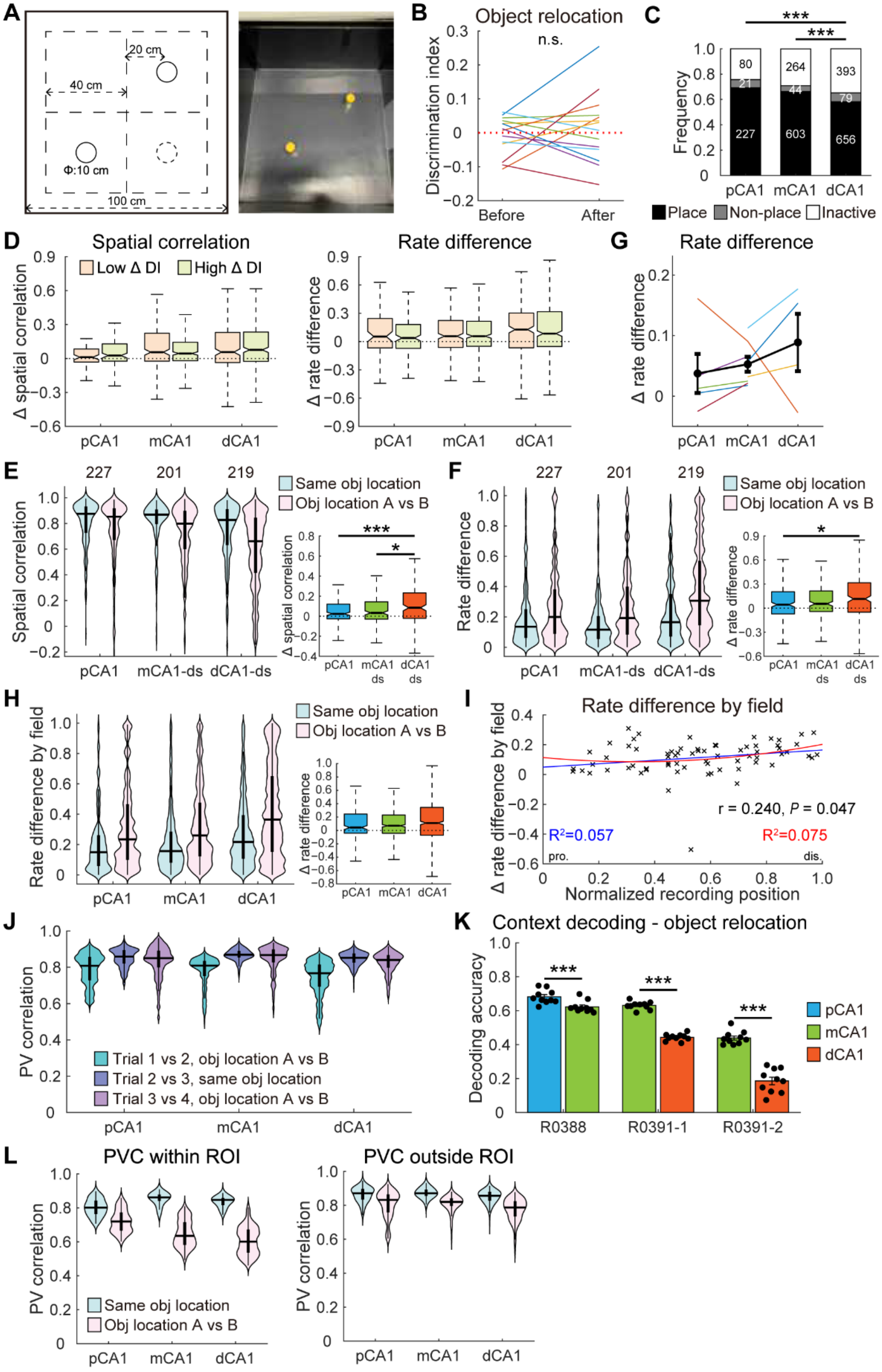

**Figure S10, related to Figure 5. Extended analyses for the object-relocation task**
**A.** Object displacement in the object-relocation task.
**B.** Discrimination index (DI) between the two objects before and after object relocation for each rat. The mean DI was close to unbiased exploration and remained largely unchanged (before: −0.007 ± 0.015; after: 0.014 ± 0.027, paired *t*-test, 14 rats, t = −0.716, *P* = 0.487). n.s., not significant.
**C.** Distribution of pyramidal neurons across three CA1 bands in the task. Proportion of active cells: pCA1, 75.61%, mCA1, 71.02%, and dCA1: 65.16%; chi-square test, $\chi^2$ = 16.157, *P* < 0.001; Holm-Bonferroni *post hoc* tests, pCA1 vs mCA1, *P* = 0.343; mCA1 vs dCA1, *P* = 0.015; pCA1 vs dCA1, *P* = 0.001. Proportion of place cells: pCA1, 69.21%, mCA1, 66.19%, and dCA1, 58.16%, $\chi^2$ = 20.656, *P* < 0.001; results of *post hoc* tests shown in the plot, ***, *P* < 0.001.
**D.** Changes in SC (left) and RD (right) between trial pairs, analyzed separately according to DI change in the object region, with outliers omitted. No difference was observed in each CA1 subregion. Mann-Whitney U tests, all *P* > 0.05.
**E**–**F.** Left: distributions of downsampled SC (**E**) and RD (**F**) in CA1 subregions in the task. Numbers on top indicate cell number after downsampling. Similar results were obtained with downsampled data. SC: two-way repeated measures ANOVA, relocation: F (1, 644) = 98.022, *P* < 0.001; band: F (2, 644) = 17.428, *P* < 0.001; relocation × band: F (2, 644) = 6.236, *P* = 0.002. RD: relocation: F (1, 644) = 108.721, *P* < 0.001; band: F (2, 644) = 14.763, *P* < 0.001; relocation × band: F (2, 644) = 3.018, *P* = 0.050. Right: changes in activity measures between condition pairs based on downsampled data, with outliers omitted. SC: pCA1, 0.024 (−0.026–0.121), mCA1, 0.033 (−0.030–0.144), and dCA1, 0.085 (−0.021–0.232); Kruskal-Wallis test, 647 cells, H = 15.013, *P* = 0.001. RD: pCA1, 0.044 (−0.072–0.204), mCA1, 0.055 (−0.042–0.215), and dCA1, 0.116 (−0.050–0. 317); H = 7.038, *P* = 0.030. Holm-Bonferroni *post hoc* tests, *: *P* < 0.05; ***, *P* < 0.001.
**G.** ΔRD between trial pairs showed a trend of increase from pCA1 to dCA1. Data from the 8 rats with simultaneous recordings from more than one CA1 subregion. Color traces indicate median data of individual animals; black trace represents mean ± SEM across animals. Rat R0172 showed a reversed trend, possibly due to low cell numbers in each subregion (26, 16 and 13 cells, respectively).
**H.** Left: distribution of $RD_f$ across CA1 subregions between trials with relocated objects (Same obj location) and trials before and after object relocation (Obj location A vs B) in the task. Two-way repeated measures ANOVA, relocation: F (1, 1 483) = 183.183, *P* < 0.001; band: F (2, 1 483) = 26.108, *P* < 0.001; relocation × band: F (2, 1 483) = 1.724, *P* = 0.179. Right: degree of changes in $RD_f$ between condition pairs, with outliers omitted. pCA1: 0.043 (−0.040–0.246), mCA1: 0.070 (−0.041–0.231), and dCA1: 0.111 (−0.072–0.344); Kruskal-Wallis test, 1 486 cells, H = 4.467, *P* = 0.107.
**I.** Correlation between normalized recording position and changes in $RD_f$ between condition pairs. Each cross corresponds to average data of a single tetrode. Blue curve, linear regression; red curve, quadratic polynomial regression.
**J.** Distribution of overall PVCs of CA1 subregions between neighboring trials in the object-relocation task. A notable hysteresis effect was observed between trials 3 and 4. Two-way repeated-measures ANOVA for trial pairs with objects in the same location (Trials 2 vs 3), and trial pairs in different locations (Trials 1 vs 2), relocation: F (1, 1 197) = 1 097.785, *P* < 0.001; band: F (2, 1 197) = 20.565, *P* < 0.001; relocation × band: F (2, 1 197) = 26.839, *P* < 0.001.
**K.** Context decoding accuracy (mean ± SEM) for simultaneously recorded pCA1/mCA1 and mCA1/dCA1 data sets. Black dots represent data from individual predictions. The three experiments with more than 30 simultaneously recorded place cells in both bands are shown. Downsampling was applied to equalize cell numbers. R0388: pCA1, 0.682 ± 0.013, mCA1, 0.623

± 0.011; paired *t*-test, t = 4.731, $P = 0.001$, 10 calculations. R0391-1: mCA1, 0.631 ± 0.007, dCA1, 0.443 ± 0.006; t = 27.39, $P < 0.001$. R0391-2: mCA1, 0.439 ± 0.012, dCA1, 0.186 ± 0.022; t = 10.44, $P < 0.001$. ***: $P \leq 0.001$.

**L.** Correlations of PVs within (left panel) and outside (right panel) ROI, between trial pairs with objects in the same location and trial pairs in different locations. PVC within ROI: two-way repeated measures ANOVA, relocation: $F (1, 237) = 1556.030$, $P < 0.001$; band: $F (2, 237) = 13.251$, $P < 0.001$; relocation × band: $F (2, 237) = 108.992$, $P < 0.001$. PVC outside ROI: relocation: $F (1, 957) = 1085.693$, $P < 0.001$; band: $F (2, 957) = 37.345$, $P < 0.001$; relocation × band: $F (2, 957) = 7.099$, $P = 0.001$.

**Fig. S11**

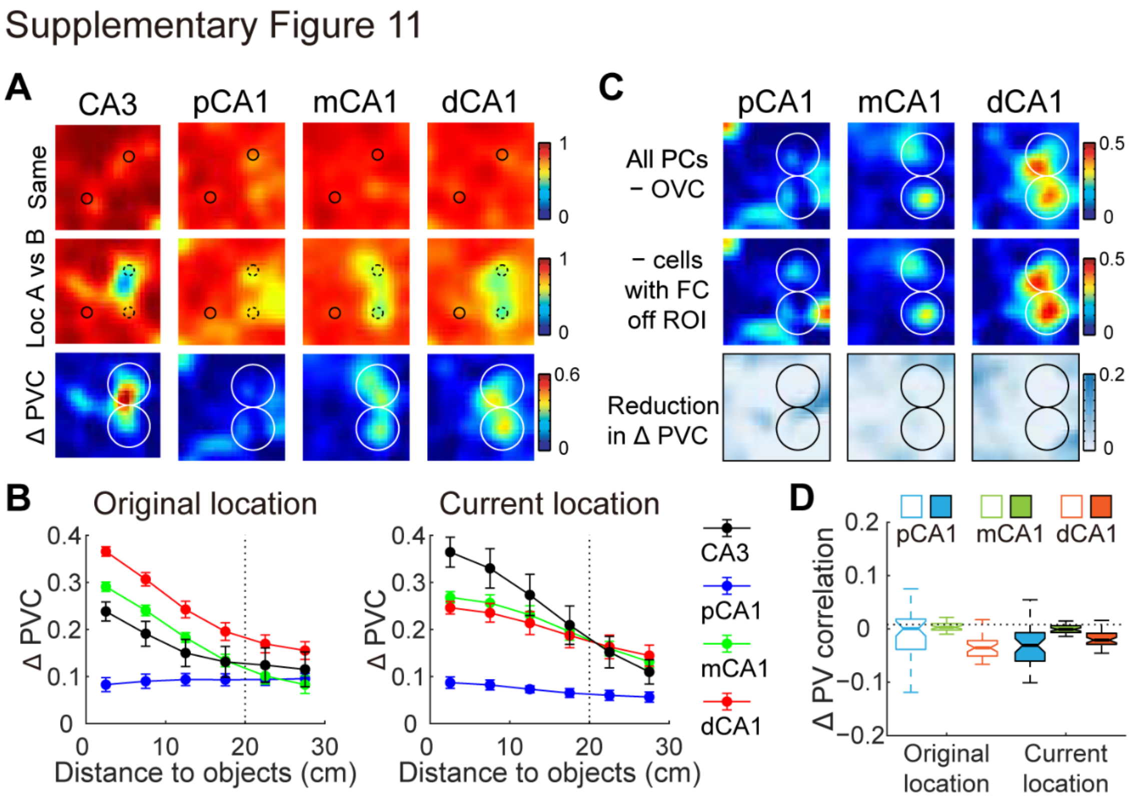


**Figure S11, related to Figure 6. Extended analyses for CA3 and CA1 responses to the displaced object**

**A.** Heatmaps showing PVC of all putative pyramidal neurons in each CA1 subregion, as well as upstream CA3, between trials (top two columns), together with changes in PVC between trial pairs (bottom column) in the object-relocation task. Small circles represent the objects; large circle indicates ROI.

**B.** Degree of changes in PVC as a function of distance to the relocated object (mean ± SEM) at the original location (left) and the current location (right) in each CA1 subregion, as well as upstream CA3 (259 neurons). CA3 showed significant activity changes at the current location, but a less prominent response at the original location, Mann-Whitney U test, 168 bins, Z = 2.776, $P$ = 0.006. PVC changes across subregions: original location: two-way ANOVA, distance: F (5, 312) = 10.324, $P < 0.001$; band: F (3, 312) = 24.689, $P < 0.001$; distance × band: F (15, 312) = 1.687, $P$ = 0.052. Holm-Bonferroni *post hoc* tests, dCA1 showed the highest PVC changes, all $P < 0.001$; CA3 and mCA1 exhibited a higher PVC change compared to pCA1, but a lower change than dCA1, all $P \leq 0.003$. Current location: distance: F (5, 312) = 13.297, $P < 0.001$; band: F (3, 312) = 36.053, $P < 0.001$; distance × band: F (15, 312) = 1.666, $P$ = 0.056. *post hoc* tests, pCA1 displayed the lowest PVC changes, all $P < 0.001$.

**C.** Heatmaps showing changes in PVC for population of place cells excluding cells with field changes outside ROI, or cells displaying rate changes within receptive fields, as well as reduction of PVC compared to population response without OVC-like neurons. Small circles represent objects, large circles indicate ROI. PC: place cell; FC: field change.

**D.** Reductions in PVC at original and current object locations after exclusion of cells with field changes outside ROI, with outliers omitted. Removing these neurons from analysis had minimal influence to population responses within ROI. Original location: pCA1, 0.001 (−0.038–0.018), mCA1, 0.003 (−0.002–0.010), and dCA1, −0.036 (−0.051–−0.022). Current location: pCA1, −0.031 (−0.060–−0.007), mCA1, −0.001 (−0.007–0.006), and dCA1, −0.021 (−0.029–−0.008).

**Fig. S12**

Supplementary Figure 12

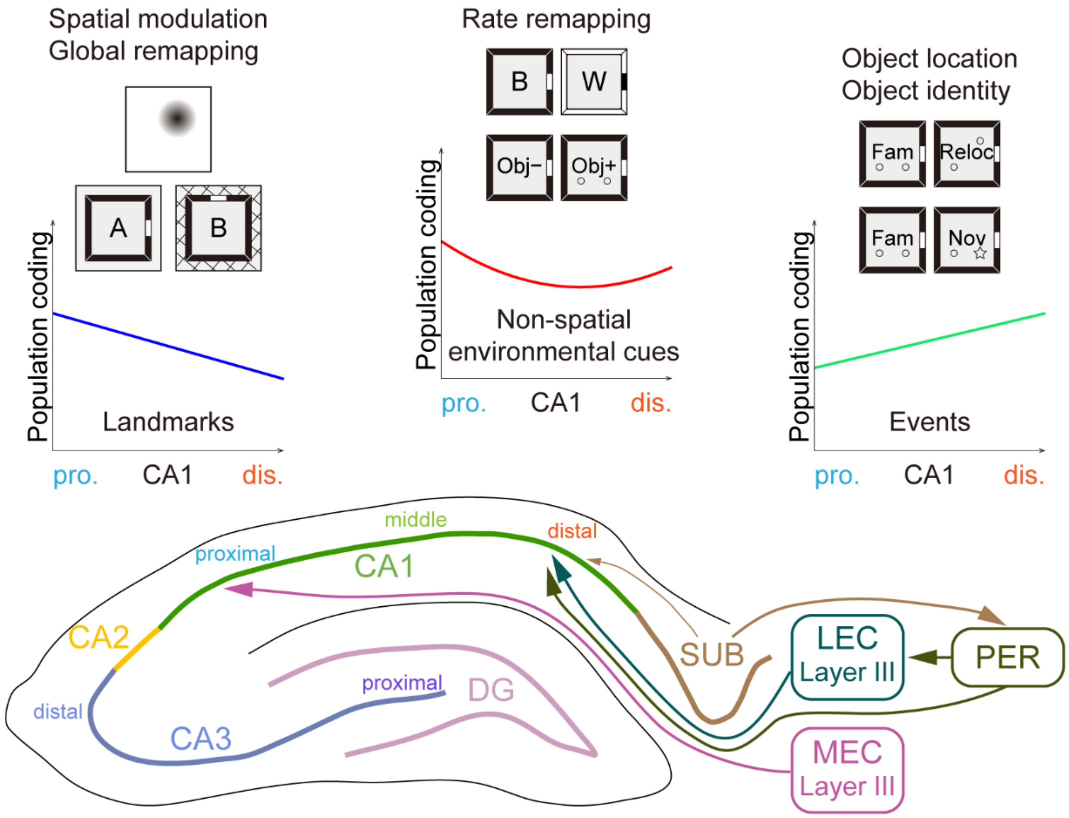


**Figure S12, related to discussion. Summary of mnemonic functions and related afferent connections that varied along CA1 transverse axis**

The three distinct patterns of population coding that differ along CA1 proximodistal axis. The medial entorhinal cortex is responsible for processing information related to navigation, such as direction and distance to landmarks, position, borders and speed. In contrast, the lateral entorhinal cortex supplies information pertinent to ongoing experience, including egocentric object vectors, object identity, memory traces, odors and time.
